# Climate-driven specialisation in plant–pollinator networks peaks outside the tropics

**DOI:** 10.1101/2025.10.08.680666

**Authors:** Sailee P. Sakhalkar, Nico Blüthgen, Laura A. Burkle, Paul CaraDonna, Bo Dalsgaard, Carsten F. Dormann, Christopher N. Kaiser-Bunbury, Tiffany M. Knight, Jeff Ollerton, Julian Resasco, Matthias Schleuning, Diego P. Vázquez, Rita N. Afagwu, Ruben Alarcón, Felipe W. Amorim, Marsal D. Amorim, Dominik Anýž, Blanca Arroyo-Correa, Maddi Artamendi, Zeynep Atalay, Justin A. Bain, Katherine C. R. Baldock, Gavin Ballantyne, Caio S. Ballarin, Behnaz Balmaki, Michael Bartoš, Vasuki Belavadi, Paolo Biella, Anne D. Bjorkman, João Paulo Raimundo Borges, Jordi Bosch, Camila Bosenbecker, Stephane Boyer, Karin T. Burghardt, Edson Cardona, Quebin Casiá, Anthony M. T. Castagna, Ferhat Celep, Melanie N. Chisté, Yann Clough, James M. Cook, Liam P. Crowther, Fabiana D. Reis, Luis P. da Silva, Wesley Dáttilo, Valeska De Cárdenas, Joan Díaz-Calafat, Lynn V. Dicks, Rye Dickson, Marion L. Donald, Lee A. Dyer, Fairo F. Dzekashu, Natalia Escobedo Kenefic, Tom M. Fayle, Laura L. Figueroa, Jan Filip, Raúl García-Camacho, Amy-Marie Gilpin, Luis Giménez-Benavides, Ingrid N. Gomes, Heather Grab, Ingo Grass, Travis J. Guy, Veronica Hederström, Carlos Hernández-Castellano, Sandra Hervías-Parejo, Andrea Holzschuh, Sebastian Hopfenmüller, José M. Iriondo, Štěpán Janeček, Jana Jersáková, Shalene Jha, Aphrodite Kantsa, Tamar Keasar, Liam Kendall, Yannick Klomberg, Ishmeal N. Kobe, Theresia Krausl, Patricia Landaverde, Carlos Lara-Romero, H. Michael G. Lattorff, Felipe Librán-Embid, Tial C. Ling, Sissi Lozada-Gobilard, Francis Ewome Luma, Pedro Luna, Ludmilla M. S. Aguiar, Ana Carolina Pereira Machado, Isabel C. Machado, Ainhoa Magrach, Fabienne Maihoff, Gabriel Marcacci, Carlos Martínez-Núñez, Pietro K. Maruyama, Natsuki Matsubara, Maggie Mayberry, Marco A. R. Mello, Ugo Mendes Diniz, Marcos Méndez, Jan E. J. Mertens, Rubén Milla, Javier Morente-López, Tarcila L. Nadia, Georgios Nakas, Anders Nielsen, Alon Ornai, Sergio Osorio-Canadas, Wilhelm H. A. Osterman, Todd M. Palmer, Theodora Petanidou, Christian W. W. Pirk, Kit S. Prendergast, Richard Primack, Luis M. Primo, Marina Querejeta, Zelma Quirino, André Rodrigo Rech, Sara Reverté, Pedro J. Rey, Léo C. Rocha-Filho, Anselm Rodrigo Domínguez, Masoud A. Rostami, Avery L. Russell, Silvia Santamaría, Francisco A. R. Santos, Takehiro Sasaki, Manu E. Saunders, Victor H. D. Silva, Michael P. Simanonok, Antigoni Sounapoglou, Ingolf Steffan-Dewenter, Sam Tarrant, Rubén Torices, Anna Traveset, Teja Tscharntke, Guillermo Uceda-Gómez, Katherine R. Urban-Mead, Casper J. van der Kooi, Catrin Westphal, Abdullahi Yusuf, Robert Tropek

## Abstract

Pollination is a key ecological process sustaining biodiversity and food security, yet global patterns of plant–pollinator specialisation have remained unresolved. Using the largest global dataset of quantitative networks (>3,400 networks, >110,000 interactions), we show that the latitudinal specialisation gradient (LSG) exists, but it is non-linear, hemispherically asymmetric, and strongly taxon-dependent. Network-level and pollinator specialisation were lowest in the tropics and peaked at northern mid-latitudes, whereas plants tended to become more specialised toward higher latitudes. Climate consistently outperformed latitude, species richness, and environmental productivity as a predictor of these patterns. Specialisation declined with increasing temperature, rose with moderate rainfall before declining at the wettest sites, and increased with temperature seasonality, but plants and pollinators responded differently to these drivers. Functional groups diverged strongly: ectothermic insects were most specialised in cooler, seasonal climates, while birds showed weaker links to latitude but reduced specialisation in wetter regions. These findings demonstrate that climate, rather than latitude or species richness, structures global variation in specialisation. Because warmer and less seasonal climates promote generalisation, climate change is likely to disrupt the most specialised pollination systems, unevenly across taxa and regions, with important consequences for biodiversity and ecosystem stability.

## Introduction

Specialisation of plant–pollinator interactions fundamentally influences pollination efficiency, plant reproduction, and ecosystem functioning and stability. The extent to which species rely on limited partners shapes resource partitioning and coexistence, thereby structuring communities, affecting species persistence, and driving evolutionary processes that generate and maintain biodiversity (Armbruster & Muchhala 2009; Ollerton 2017; Van Der Niet & Johnson 2012). Specialised pollination is also hypothesised to facilitate reproductive isolation and speciation, promoting niche differentiation, and reducing competition, thus contributing to the exceptional diversity of plants and pollinators (Kay & Sargent 2009; Van Der Niet & Johnson 2012). Yet, despite its ecological and evolutionary importance, broad-scale patterns in interaction specialisation remain unresolved.

Latitudinal Specialisation Gradient (LSG), a systematic geographic pattern in the degree of interaction specialisation, remains a fundamentally unresolved question in ecology (Hargreaves 2024; Pinheiro *et al*. 2023). Clarifying whether and why LSG exists is central to understanding the origin and maintenance of biodiversity gradients, community assembly, and the evolution of species interactions (Moles & Ollerton 2016; Schleuning *et al*. 2012). LSG is closely linked to the well-established latitudinal diversity gradient (LDG), which describes increasing species richness toward the equator (Armbruster & Muchhala 2009; Dobzhansky 1950; Pauw 2013; Schemske *et al*. 2009). Because specialisation and species richness are theoretically coupled, testing the existence of LSG and revealing its patterns and drivers is essential for our understanding of mechanisms structuring global biodiversity and ecosystem functioning, as well as their responses to global change (Brown 2014; MacArthur 1972; Schemske *et al*. 2009). Pollination networks are ideal for testing LSG, because plant–pollinator interactions occur across all terrestrial biomes, encompass broad phylogenetic and functional diversity, and represent mutualisms integral to community assembly and diversification (Schemske *et al*. 2009; Schleuning *et al*. 2012; Vázquez & Stevens 2004).

Theoretical predictions and empirical evidence for LSG conflict sharply. Two prominent theoretical frameworks predict stronger specialisation in the tropics (Hargreaves 2024; Schemske *et al*. 2009): the latitude–niche breadth hypothesis, attributing narrower niches to competition and benign climates (MacArthur 1972), and the biotic interactions hypothesis, linking co-evolutionary specialisation to long-term climatic stability (Dobzhansky 1950). Supporting empirical evidence for plant-pollinator LSG includes increased specialisation in plant-hummingbird networks toward the equator (Dalsgaard *et al*. 2011) and a global synthesis of 54 pollination networks reporting higher specialisation at lower latitudes (Trøjelsgaard & Olesen 2013). In direct contrast, optimal foraging theory predicts more generalised interactions in the tropics because high diversity often coincides with low local abundances of individual interacting species (Schleuning *et al*. 2012; Vázquez & Stevens 2004). This prediction is consistent with a global analysis of 58 pollination networks showing decreasing network-level specialisation toward the tropics (Schleuning *et al*. 2012). Further, Moles and Ollerton (2016) argued against any universal LSG, proposing that specialisation patterns vary with interaction type, interacting taxa, and ecosystem context, as supported by studies finding no consistent LSG (Luna *et al*. 2022; e.g. Ollerton & Cranmer 2002; Rahimi & Jung 2024).

Much of the variation in the existing studies can be attributed to geographic, taxonomic, and methodological biases. The global analyses have been dominated by temperate networks from the Americas and Europe, leaving critical gaps elsewhere (Ollerton 2017; Vizentin-Bugoni *et al*. 2018), and have often ignored regional differences, despite the described strong ecosystem-specific effects on specialisation (Pauw & Stanway 2015). Many studies examined a single group (bees in Cirtwill *et al*. 2025; e.g. hummingbirds in Dalsgaard *et al*. 2011; cacti in Gorostiague *et al*. 2023) or combined groups indiscriminately (Luna *et al*. 2022; e.g., Olesen & Jordano 2002; Trøjelsgaard & Olesen 2013), potentially masking group-specific patterns shaped by contrasting diversity gradients, evolutionary histories, and resource-use strategies (Armbruster 2017; Ollerton *et al*. 2007). Methodologically, cross-study comparisons are further hampered by the use of non-standardised specialisation metrics, uneven sampling effort, and the mixing of complete community networks with taxon-constrained partial networks (Armbruster 2017; Brimacombe *et al*. 2022; Doré *et al*. 2021; Ollerton *et al*. 2007; Vizentin-Bugoni *et al*. 2018). Overcoming these biases is essential for the rigorous testing of plant-pollinator LSG.

Here, we assemble the most comprehensive global dataset of quantitative plant– pollinator networks to date (Fig. 1), comprising 3,415 networks from 162 studies, covering 110,571 pairwise interactions among 5,343 pollinator and 6,126 plant species, spanning all major terrestrial biomes across a broad latitudinal range (Fig. 2). We address the limitations of prior syntheses by standardising sampling completeness, applying robust specialisation metrics (network-level *H*_*2*_*’* and species-level *d’*), and accounting for methodological differences. Crucially, we analyse LSGs separately for plants and functional pollinator groups, testing predictions that latitudinal patterns reflect their ecological and evolutionary differences. Finally, we test whether specialisation is driven by plant and pollinator diversity, climate, or environmental productivity, providing deeper insight into processes structuring global variation in plant–pollinator specialisation (Fig. 1). By overcoming these geographic, taxonomic, and methodological constraints and explicitly analysing functional-group-specific patterns, our study advances ecological theory on biodiversity patterns, evolutionary ecology, and the architecture of mutualistic networks.

**Figure 1.**
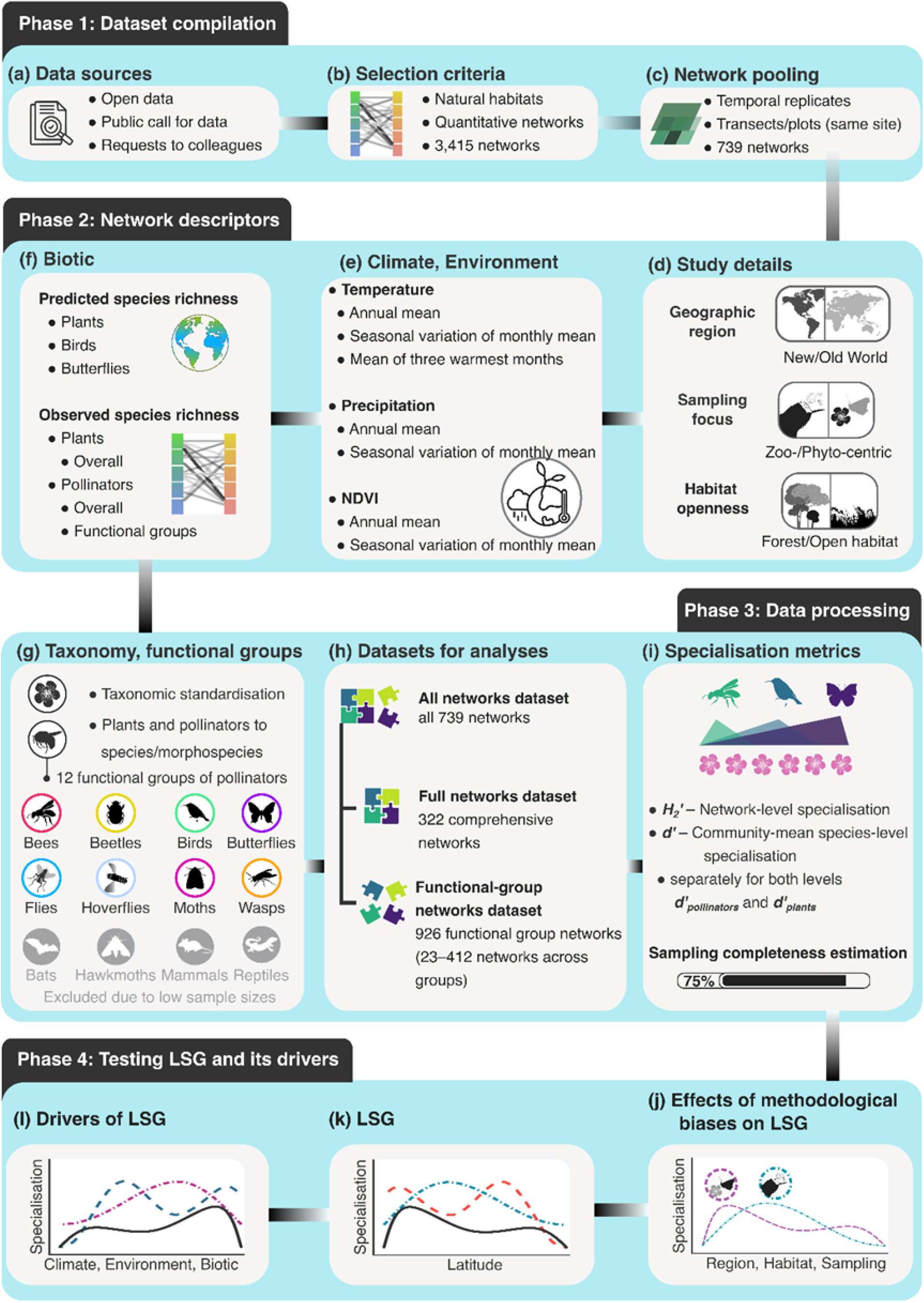
Conceptual workflow for analysing the latitudinal specialisation gradient (LSG) in plant-pollinator interactions. **Phase 1:** (a) compilation of available plant–pollinator networks, (b) application of predefined selection criteria, and (c) pooling of spatiotemporal network replicates from the same site to minimize non-focal variation. **Phase 2:** (d–f) characterisation of networks with 28 descriptors (Table S2). **Phase 3:** (g) taxonomic standardisation and assignment of pollinators to functional groups (grey icons indicate groups excluded from functional-group analyses due to data insufficiency), followed by (h) construction of three complementary datasets and (i) calculation of three specialisation metrics and sampling completeness for each sampling network. **Phase 4:** (j–l) multi-stage modelling of LSG, including (j) evaluation of potential methodological biases, (k) fitting latitudinal models with significant biases as random effects, and (m) testing climatic, environmental, and biotic predictors to identify LSG drivers. See Online Methods for details.

**Figure 2.**
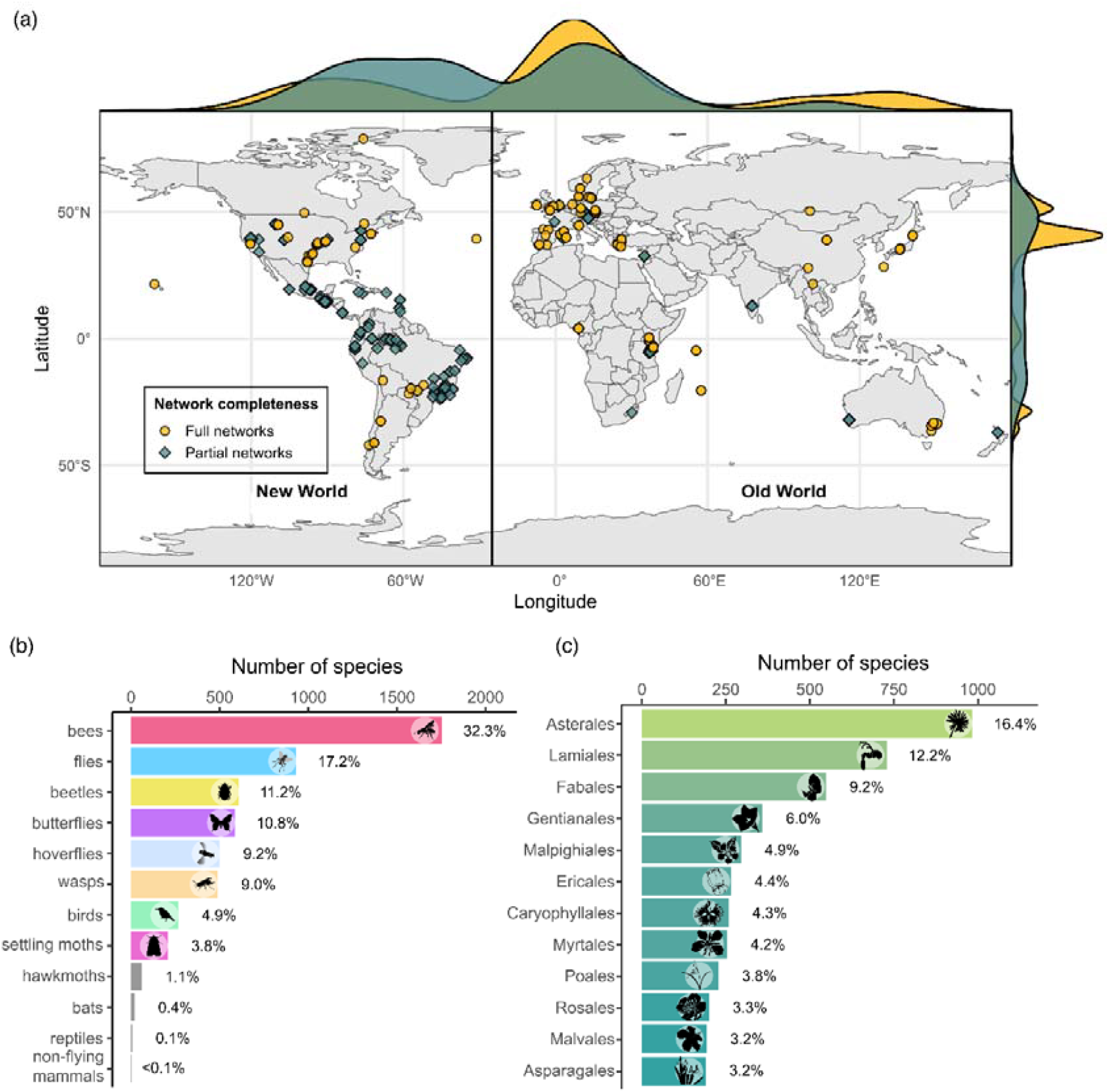
Global coverage and taxonomic composition of the analysed plant–pollinator networks. (a) Geographic distribution of networks, grouped into the New World and Old World. Symbols denote network completeness: taxonomically restricted (partial) and unrestricted (full). Network latitudinal ranges per region are indicated below the map. Marginal density plots show the distributions of network locations by their completeness along latitude (right) and longitude (top). (b) Pollinator composition by functional group, with bars indicating the percentage of all pollinator species in the dataset. (c) Plant composition, showing the 12 most speciose orders and their percentages in the dataset.

## Results

### Latitudinal specialisation gradient (LSG)

We first tested whether interaction specialisation follows systematic latitudinal patterns, directly addressing the LSG hypothesis (Fig. 1j–k). Using generalised additive mixed models (GAMMs) and hierarchical GAMMs (HGAMs) on the three datasets (all networks, full networks, and functional-group networks), we modelled the three specialisation metrics (network-level *H*_*2*_*’*, and community-mean species-level *d’*_*plants*_ and *d’*_*pollinators*_) against latitude, while accounting for potential methodological biases of the general patterns from previous studies (Fig. 3; Tables S4–S5). Geographic region and habitat openness most frequently showed significant effects (Table S4; Fig. S1) and were therefore retained as random effects in the final models (Table S5).

**Figure 3.**
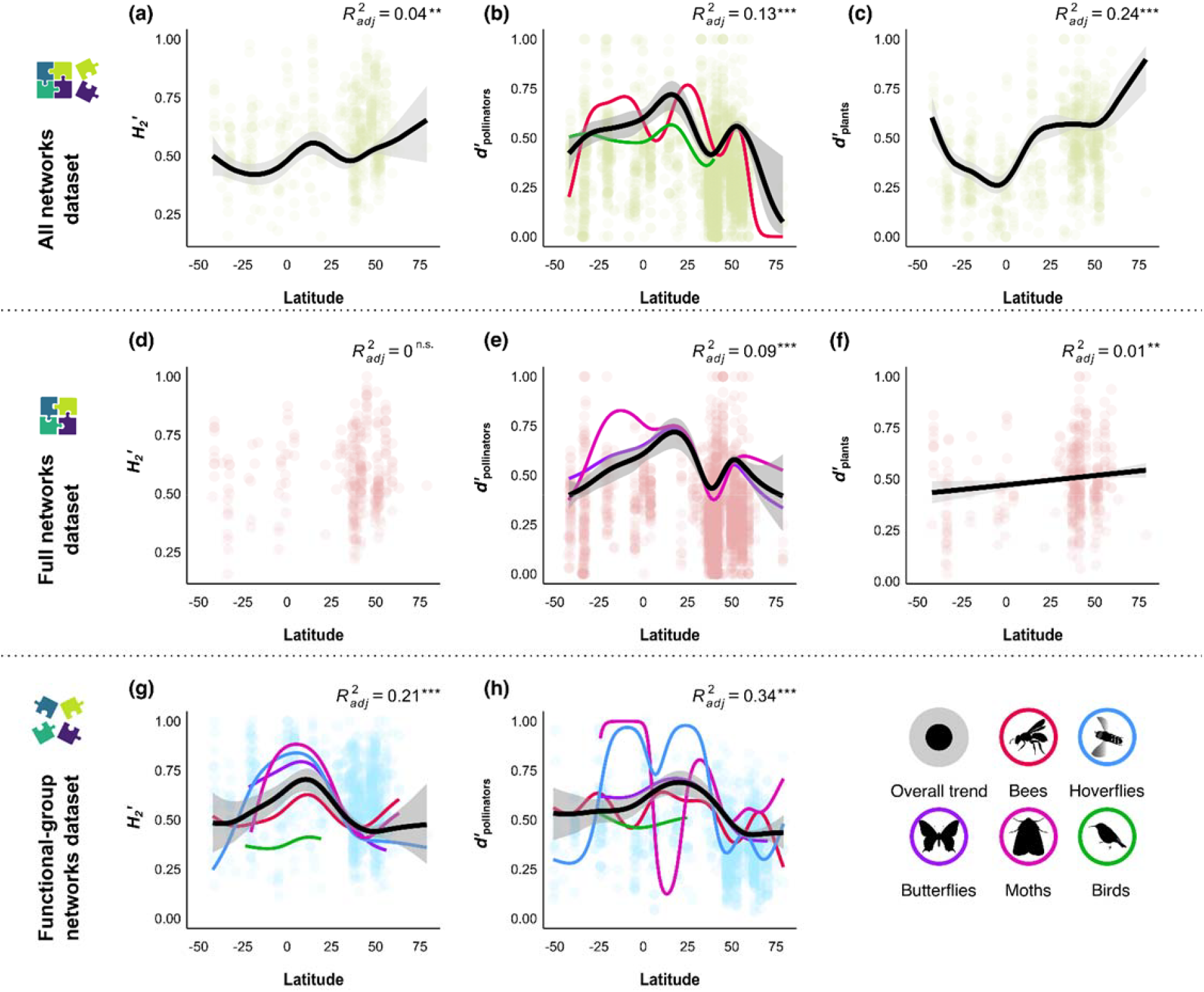
Latitudinal patterns in specialisation of plant–pollinator interactions across datasets and metrics. Rows show datasets: (a–c) all networks; (d–f) full networks; (g–h) functional-group networks. Columns show specialisation metrics: network-level *H*_*2*_*’* (left); and community-mean species-level separated for pollinators (*d’*_*pollinators*_, middle) and for plants (*d’*_*plants*_, right; not calculated for the functional-group networks dataset). Curves are GAMM/HGAM fitted smooths: black curves indicate overall latitudinal trends with 95% confidence intervals (grey shading), coloured curves represent significant functional group-specific trends (see legend). Coloured points are raw data. Curves are drawn only when the smooth term for latitude is significant; otherwise only points are shown. Adjusted *R*^*2*^ and significance (**p < 0.01; ***p < 0.001; n.s.: not significant) are reported in each panel, model summaries are reported in Table S5.

Across models, latitudinal patterns in specialisation were generally non-linear and hemispherically asymmetric, with several models showing a mid-latitude maximum around ∼25–40°N (Fig. 3). Latitude explained little variation in the all networks and full networks datasets (*R*^*2*^ ≤ 0.13), but resolving networks by functional groups consistently increased explanatory power of the models (functional-group networks dataset: *R*^*2*^ = 0.21–0.34; *d’*_*plants*_ in the all networks dataset: *R*^*2*^ = 0.24) and revealed taxon-dependent LSGs (Table S5; Fig. 3).

In the all networks dataset, *H*_*2*_*’* and *d’*_*plants*_ showed a nearly U-shaped pattern, lowest in the tropics and increased toward higher latitudes, especially in the Northern Hemisphere (Fig. 3a,c). *d’*_*pollinators*_ was bimodal, with maxima in the northern subtropical and temperate latitudes (Fig. 3b; Table S5). Among functional pollinator groups, only bees and birds exhibited significant *d’* patterns (Table S5): bees followed a generally hump-shaped pattern with dips in the tropics and northern temperate latitudes, whereas birds showed an almost flat pattern with an apparent dip in the northern temperate zone (Fig 3b). In the full-network dataset, latitude significantly predicted both *d’* metrics but not *H*_*2*_*’* (Fig. 3d–f; Table S5). *d’*_*pollinators*_ followed a broadly hump-shaped pattern peaking in the northern subtropics, with more complex variation at higher northern latitudes; butterflies and moths broadly mirrored this pattern, although moths showed an additional peak in the southern tropics (Fig. 3e). *d’*_*plants*_ increased approximately linearly from the south to the north but with very low explained variation (Fig. 3f). In the functional-group networks dataset, latitude significantly influenced both *H*_*2*_*’* and *d’*_*pollinators*_ (Fig. 3g–h; Table S5). *H*_*2*_*’* followed a broadly U-shaped pattern with highest values in the northern subtropics, particularly for butterflies, moths, hoverflies, and bees, although some of these groups also exhibited mild secondary rises toward the lowest and highest latitudes (Fig. 3g). *d’*_*pollinators*_ showed a broadly hump-shaped pattern peaking at northern mid-latitudes and declining slightly toward the tropics and strongly toward the northern subarctic (Fig. 3h). Nevertheless, functional pollinator groups deviated from this pattern: butterflies generally mirrored it, whereas moths, hoverflies, bees, and birds showed irregular responses, mostly with multiple maxima and minima (Fig. 3h).

### Drivers of LSG

We next tested whether forward selected climatic (mean annual temperature, MAT; mean annual precipitation, MAP; and temperature seasonality, SD_MMT), environmental (NDVI and SD_NDVI), and biotic (12 measures of predicted and observed species richness of plants and pollinators) variables explained global variation in specialisation (*H*_*2*_*’, d’*_*pollinators*_, and *d’*_*plants*_). Using the same GAMM/HGAM framework (Fig. 1i), we compared individual predictors (and the MAT×MAP interaction) among each other and against the corresponding latitudinal models (Fig. 4; Tables S6 and S7). All three potential methodological biases of the general patterns (geographic region, sampling focus, and habitat openness) showed significant effects in some models (Table S4) and were therefore retained as random effects where applicable.

**Figure 4.**
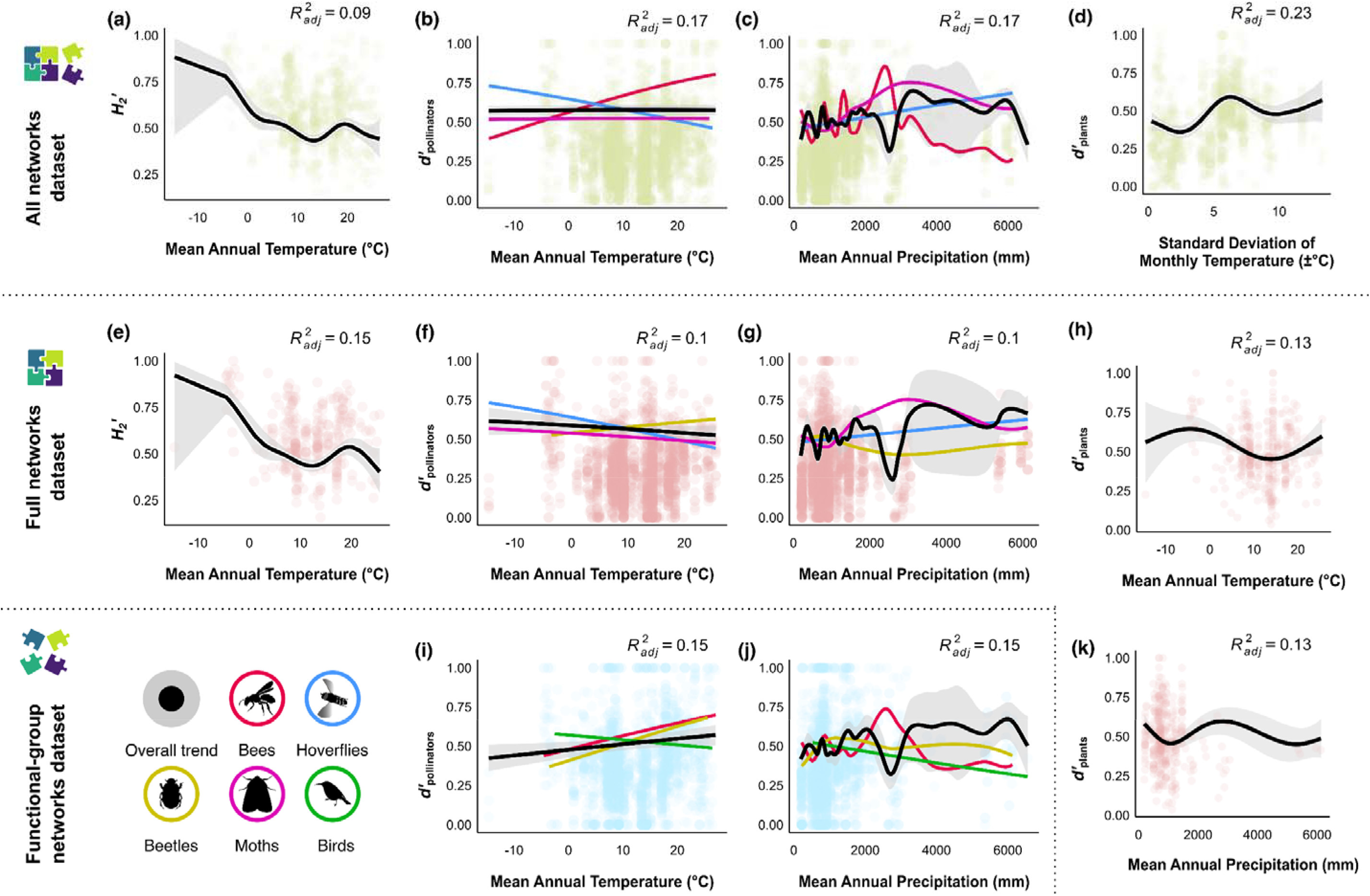
Most important climatic predictors for plant–pollinator specialisation across datasets and metrics. Panels show, for each metric–dataset combination, the predictors from the best-supported GAMM/HGAM model (lowest ΔAIC relative to the latitude-only baseline; see Table S7). Rows broadly correspond to datasets: (a–d) all networks; (e–h, k) full networks; (i–j) functional-group networks. Columns present specialisation metrics: network-level *H*_*2*_*’* (left); and community-mean species-level separated for pollinators (*d’*_*pollinators*_, middle) and for plants (*d’*_*plants*_, right; not calculated for the functional-group networks dataset). Curves are GAMM/HGAM fitted smooths: black curves indicate overall latitudinal trends with 95% confidence intervals (grey shading), coloured curves represent significant functional group-specific trends (see legend). Coloured points are raw data. The functional-group networks dataset model for *H*_*2*_*’* is omitted because latitude was the best predictor and is shown in Fig. 3g. Adjusted R^2^ is reported in each panel; model details are in Table S7.

Across models, predictor–response relationships were frequently non-linear. Except for *H*_*2*_*’* in the functional-group networks dataset, where latitude remained the strongest predictor (Fig. 3g), climatic variables (especially MAT, MAP, or their interaction) most consistently explained variation in interaction specialisation, whereas environmental and biotic predictors never did (Fig. 4). However, even the best climate-based models often explained less variance than the corresponding latitudinal models (Tables S6–S7).

For *H*_*2*_*’*, MAT was the best predictor in the all networks and full networks datasets, showing a non-linear but generally declining relationship with increasing temperature (Fig. 4a,e; Tables S6–S7). For *d’*_*pollinators*_, MAT×MAP consistently provided the best-fitting models across all datasets (Fig. 4b,c,f,g,I,j). *d’*_*pollinators*_ responses to MAT were linear but varied in direction: increasing in the all networks and functional-group networks datasets but declining in the full network dataset. Functional groups diverged further, with bees and beetles generally increasing in specialisation with MAT, hoverflies declining, and other groups showing less consistent responses. In contrast, MAP effects were more uniform, with *d’*_*pollinators*_ generally increasing with precipitation but dipping near 3000 mm annually (Fig. 4c,j,g). For *d’*_*plants*_, SD_MMT was the strongest predictor in the all networks dataset, with specialisation generally increasing under greater temperature seasonality but showing local minima around ±3°C and ±10°C (Fig. 4d). MAT×MAP was the most plausible predictor of *d’*_*plants*_ in the full networks dataset, with an S-shaped response to MAT dipping just below 0°C and peaking near 12°C (Fig. 4h), and a nearly hump-shaped relationship with MAP with the highest values at ∼3000 mm annually (Fig. 4k).

## Discussion

Our global analyses of >3,400 quantitative plant–pollinator networks provide the strongest evidence to date that the latitudinal specialisation gradient (LSG) exists, but it is neither simple nor universal. Latitudinal patterns in specialisation were non-linear, differed between hemispheres, and varied across taxa. Most importantly, specialisation of plant–pollinator interactions did not peak in the tropics: plants tended to be more specialised toward higher latitudes, network-level and pollinator specialisation peaked in northern subtropical and temperate regions, and functional pollinator groups partly diverged in responses. These patterns held after accounting for methodological biases (Moles & Ollerton 2016; e.g. Vázquez *et al*. 2009), and resolving pollinators by functional group substantially increased the variation explained by latitude. Across the specialisation metrics and datasets, climate consistently provided stronger explanatory power than latitude or diversity, with temperature, precipitation, and their interaction emerging as the dominant drivers of global variation in specialisation, while environmental productivity and species richness played no significant role.

None of the main theoretical frameworks fully explain the revealed global patterns in plant–pollinator specialisation. The latitude–niche breadth (MacArthur 1972) and biotic interactions (Dobzhansky 1950) hypotheses both predict highest specialisation in the tropics, yet our analyses revealed both plants and pollinators to be most generalised at tropical latitudes. This pattern is consistent with the optimal foraging theory, which predicts that species-rich tropical communities favour generalisation because high diversity reduces per-species abundance and encounter rates, thereby promoting broader diets (Schleuning *et al*. 2012; Vázquez & Stevens 2004). The additional declines in pollinator specialisation toward the highest latitudes may reflect physiological constraints in extreme climates. We lacked data on pollinator abundance and floral rewards, the resources for plants and pollinators respectively, to test the proposed mechanisms of optimal foraging theory directly. Nevertheless, because neither species richness nor environmental productivity (a potential proxy for resource availability; Bailey *et al*. 2004; Pauw 2013) outperformed climate predictors in our models, these mechanisms are unlikely to underlie the observed LSG patterns (Hargreaves 2024; MacArthur 1972; Schemske *et al*. 2009; Vázquez & Stevens 2004).

Climate emerged as the primary driver of the global variation in pollination specialisation. The declining network-level and pollinator specialisation towards warmer environments confirms that the predicted simple latitudinal gradients are in fact modified by elevation and regional context. These patterns also contradict expectations that warmer, less seasonal ecosystems foster higher specialisation in plant–pollinator interactions (Granot & Belmaker 2020; Petanidou *et al*. 2018; Schemske *et al*. 2009). Instead, in warmer environments, pollinators may extend their activity despite the known high phenological turnover of tropical plants (Du *et al*. 2024; e.g. Song *et al*. 2022), so an average pollinator species must rely on more plant species over their lifetimes or activity periods. Conversely, colder and more seasonal climates compress both flowering and foraging into shorter phenological windows, which may promote stronger specialisation (Glaum *et al*. 2021). Precipitation exerted a non-linear influence: specialisation generally increased from dry to intermediate–high rainfall but declined at the wettest sites, consistent with moderate rainfall supporting flowering and foraging, in contrast to excessive rainfall suppressing flight and diluting nectar (Aizen 2003; Klomberg *et al*. 2022; Maicher *et al*. 2018). Plant specialisation showed different climatic responses, with a broadly positive association with temperature seasonality, consistent with synchronous flowering and tighter pairings where seasonal forcing is strong (Altermatt 2010; Glaum *et al*. 2021). By showing that climate, rather than latitude or biodiversity alone, structures specialisation worldwide, we bridge decades of debate on the LSG and reveal the mechanisms underlying interaction diversity.

The functional-group differences demonstrate that LSG arises from group-specific traits related to climate. Plants became more specialised in highly seasonal ecosystems at higher latitudes, likely due to tighter flowering synchrony (Du *et al*. 2024; Rathcke & Lacey 1985), whereas pollinators responded more directly to temperature and precipitation (Dalsgaard *et al*. 2011). Several insect groups substantially contributed to the northern mid-latitude peak in specialisation, while birds showed only a shallow specialisation increase toward higher latitudes. These contrasts reflect differences in thermal physiology, where relatively colder and seasonal climates may select for more efficient specialised foraging in insects (Brown 2014; Dzekashu *et al*. 2023), whereas mostly tropical nectarivorous birds can adapt behaviourally to year-round available but phenologically shifting resources (Chmel *et al*. 2021). Weak or absent climatic trends in specialisation in some insect groups, including beetles, flies, and wasps, probably reflect their heterogeneous diets, although some clades are highly selective nectar or pollen feeders (Ollerton 2017; Willmer 2011).

Warmer and less seasonal climates promote generalisation over specialisation, with major implications for the fate of pollination networks under climate change. Projected reductions in seasonal differences in temperate regions (Adam *et al*. 2023; Xu *et al*. 2013) and intensified precipitation extremes in the subtropics (Wang *et al*. 2024) are likely to affect the most specialised plant–pollinator communities. Specialised interactions are particularly vulnerable to phenological mismatches, as even small shifts in flowering or pollinator activity can cause breakdowns, whereas generalists may adjust or rewire their interactions (Bartomeus *et al*. 2011; Kudo & Ida 2013). Resulting losses of specialised species could shift communities toward generalists, with uncertain consequences for ecosystem functioning and stability (Burkle *et al*. 2013). Responses may be expected to differ among groups: ectothermic insects, most specialised in cooler, seasonal climates, may be highly sensitive to resource shifts under climate warming, while bird specialisation is more strongly linked to precipitation and thus vulnerable to altered rainfall regimes. Geographic context also matters, since tropical networks are already relatively generalised, while more specialised temperate and subtropical communities may be more prone to species loss or phenological disruption. Overall, our results caution that climate change will not affect all plant–pollinator networks equally, underscoring the need to account for both climatic drivers and functional-group differences when assessing the future of pollination services and biodiversity.

## Acknowledgements

We are grateful to everybody who assisted with sampling of the analysed datasets, especially to Elisângela L. S. Bezerra, Kryštof Chmel, Joel Queiroz, and Pat Willmer; to Ondřej Mottl for priceless statistical advice; and to Sara D. Leonhardt, Martin Volf, Tereza Kočárková, Fotoula Papandreou, Javier Oñate-Casado, Riccardo Pernice, and Constantinos Charalambous for feedback on earlier versions of this manuscript. We used GPT large language model (ChatGPT, OpenAI, models 4.5, 4o, 4-turbo, and 5, June 2024–September 2025) to assist in improving the clarity, readability, and stylistic refinement of the manuscript text. This study was funded by the Czech Science Foundation project 21-24186M (to S.P.S., D.A., J.F., Š.J., I.K., A.S., R.T.). Individual datasets and co-authors were supported by Alexander von Humboldt Foundation (1134644), São Paulo Research Foundation (2023/03083-6, 2023/02881-6, 2023/17728-9), and Consulate General of France in São Paulo (all previous to M.A.M.); Bavarian State Ministry of Science and Art (to F.M.); Biotechnology and Biological Sciences Research Council (to L.V.D.); Center for Research on Biodiversity Dynamics and Climate Change, CEPID–FAPESP (2021/10639-5; to F.A. and C.S.B.); National Council for Scientific and Technological Development, CNPq (308559/2022-3 to F.A., C.S.B.; 141736/2020-8 to A.C.M.; 311665/2022-5, 400904/2019-5, and 423939/2021-1 to J.P.B.; 310508/2019-3 to I.M.; 309893/2023-2 to L.M.; 177005/2024-6 to V.S.; 305204/2024-6 to M.A.M); CAPES (Finance Code 001; COOPBRASS: 88887.947041/2024-00 to A.C.M.; 177005/2024-6 to V.S.; PROEX 88882.347259/2019-01 to U.M.); Brazilian Biodiversity Fund, FunBio (004/2021 to I.N.G.; 029/2022 to V.S.); Rufford Foundation (377031 to I.N.G.; 28478-1 to U.M.); Czech Science Foundation (19-14620S; T.F.); German Research Foundation DFG (152112243 to F.L.); Dirección General de Investigación, Universidad de San Carlos de Guatemala (4.8.63.2.27–2012 to E.C. and N.E.; 4.8.63.8.60–2018 and 4.8.63.4.41–2020 to Pa.L.); FAPEMIG (RED-00039-23 to P.M. and A.R.R.); INCT Pollination (CNPq/CAPES/FAPERJ Call 58/2022 to P.M. and A.R.R.); Faculty for Future, Schlumberger Foundation (Q.C., Pa.L.); the Human Frontier Science Program (RGP023/2023 to C.v.); European Research Council ERC (101054177 to Sa.H. and A.T.; 819374 to Y.C., V.H., and Th.K.); Knut and Alice Wallenberg Foundation (KAW 2019.0202 to A.B. and W.O.); LIFE project Olivares Vivos+ (LIFE20 NAT/ES/001487 to P.R.); Missouri Department of Conservation (K02442-PI0242-022 to R.N.A., Mag.M., and A.L.R.); National Science Foundation (DGE-2244337 to L.A.D.); OAPN project 014/2009 (to Mar.M.); CONAHCYT (CBF2023-2024-216 to W.D.); Spanish Ministry of Science, Innovation and Universities (PID2021-127900NB-I00, PGC2018-098498-A-100, and RYC2021-032351-I to A.M.); Israel Ministry of Environmental Protection (grant no. 121-5-13 to Ta.K.); and funding of iDiv via the German Research Foundation DFG (FZT 118, 202548816 to T.M.K.).

## Author contributions

S.P.S. and R.T. conceived and designed the study, with substantial input from N.B. S.P.S. collated and processed the datasets, and performed the analyses under the supervision of R.T., with feedback from N.B., B.D., C.D., C.K., D.V., J.O., J.R., L.B., M.S., P.C., and T.M.K. S.P.S and R.T. prepared all visualisations, tables, and supplementary materials, with contributions from D.A., J.F., and G.U. S.P.S. and R.T. wrote the manuscript draft, with contributions from N.B., B.D., C.D., C.K., D.V., J.O., J.R., L.B., M.S., P.C., T.M.K., and T.F. All authors contributed data, reviewed the manuscript draft, and approved its submission.

## Online Methods

All analyses, including data preparation, were performed using R 4.5.0 (R Core Team 2025).

### Dataset compilation

We compiled a global dataset of plant–pollinator interaction networks (Fig. 1a) through a systematic search of Web of Science and Google Scholar between January and May 2021 (and we extensively monitored for suitable new studies until September 2024). Search strings combined terms pollin*, network*, flower visit*, plant*. All retrieved publications were manually screened for relevance, and we also examined their cited and citing references. Publicly available datasets were incorporated from the Mangal Interaction Database (Poisot *et al*. 2016), the Web of Life database (Fortuna *et al*. 2014), and a smaller set of data from the currently non-public LifeWebs project (Butterill *et al*. 2021). To maximise geographic and methodological coverage, we also contacted authors of studies without open data and requested both published and unpublished datasets from the international community of pollination ecologists (Table S1).

Datasets were included if they met the following criteria (Fig. 1b): originated from natural or semi-natural ecosystems (excluding agricultural, urban, and other anthropogenic habitats); contained at least five plant and five pollinator species; provided directly sampled interaction data (excluding compilations over broad spatial or temporal scales); and reported quantitative interaction strengths (i.e. excluding binary datasets). Coordinates, elevation, sampling details, and other metadata were obtained from publications and/or databases when available. When essential information was missing, we contacted the original authors; studies lacking such key data were excluded. Because direct measures of pollination efficiency were rarely available, we treated all floral visitors as potential pollinators and all flower visits as pollination interactions, which is standard practice in large-scale pollination network analyses. Our compiled dataset contained 3,415 quantitative networks from 162 studies, spanning 43.60°S to 81.00°N in latitude and 1 to 4,241 m in elevation, representing virtually all terrestrial biogeographic regions and major ecosystem types (Fig. 2a, Table S1).

To reduce variation irrelevant to our aims, we pooled temporally replicated networks sampled at the same sites and merged networks representing small-scale spatial units (such as transects or plots) within the same study site (Fig. 1c). However, networks from studies explicitly examining elevational or environmental gradients were retained separately, as were networks from different publications to account for methodological differences. After these steps, our dataset contained 739 unique networks.

### Network descriptors

Each plant–pollinator network was characterised using 28 descriptors (Fig. 1d-f) grouped into four categories: study and methodological details, geographic information, environmental conditions, and local species richness (Table S2; latitudinal trends in Extended Data 2).

Study and methodological details (Fig. 1d) included the source of network data, rationale for splitting datasets into separate networks (e.g. distinct sites, elevations, habitats, or vertical strata), sampling design and field methods, sampling focus (zoocentric or phytocentric), network completeness (full or partial networks), and lower-level (plants) and upper-level (pollinators) constraints (specified if sampling was taxonomically biased). Geographic information comprised country, geographic coordinates, absolute latitude, elevation, and region (New World or Old World, Fig. 1d). Environmental conditions included habitat openness (forest or open habitat). Seven climatic and environmental variables (Fig. 1e) were extracted from geographic coordinates with the *terra* package (Hijmans et al. 2025*a*). These climatic variables were mean annual temperature (MAT, °C), standard deviation of mean monthly temperature (SD_MMT, ±°C), mean temperature of the three warmest months (°C), mean annual precipitation (MAP, mm), and the standard deviation of mean monthly precipitation (SD_MMP, ±mm) (Fick & Hijmans 2017). Environmental productivity was quantified as annual mean Normalised Difference Vegetation Index (mean NDVI; Didan 2015)and the standard deviation of monthly NDVI (SD_NDVI). Species richness descriptors (Fig. 1f) included predicted local species richness for plants (Cai *et al*. 2023), birds (Jenkins *et al*. 2013), nectarivorous birds (range maps from BirdLife, filtered using an unpublished species list by B. Dalsgaard and J. Ollerton, pers. comm.), and butterflies (Daru 2024), retrieved using the *raster* package (Hijmans *et al*. 2025b). In addition, we included observed species richness directly recorded in the original network data: plants, pollinators, and total species richness, as well as richness of individual functional pollinator groups (see Table S2 for definitions).

### Taxonomy and functional groups

We standardised the taxonomy and nomenclature for plants and pollinators across all studies (Fig. 1g) using the *taxize* package (Chamberlain & Szöcs 2013) with the GBIF database (2023). Species names were verified by exact matching, unmatched or partially matched (“fuzzy”) species names were corrected to fix typos and re-matched. Taxa identified to morphospecies were retained as unique entities (called species hereinafter) within individual datasets but were not comparable across different studies.

Pollinator species were assigned to functional groups reflecting distinct floral resource use and ecological roles, following commonly accepted pollination syndromes (Willmer 2011): bees, wasps (incl. sawflies), butterflies, hawkmoths, settling moths, beetles, birds, bats, non-flying mammals, and reptiles (Fig. 1g). Flies were further separated into hoverflies, known for their relatively specialised floral associations and high dataset representation, and all other flies without further subdivision of specialised fly groups due to low abundance and generally coarse taxonomic identification in our data. Other floral visitors (such as spiders, ants, orthopterans, hemipterans) were excluded as mostly accidental floral visitors and/or unimportant pollinators (although some species can be important in unique pollination systems; Ollerton 2021). Plants were not subdivided into functional groups.

After the taxonomic standardisation and restricting the networks to the selected pollinator functional groups, the dataset comprised 110,571 pairwise interactions among 5,343 pollinator species and 6,126 plant species, spanning a broad diversity of pollination systems (Fig. 2b–c).

### Datasets for analyses

To reflect the dataset heterogeneity and robustly test latitudinal specialisation patterns, we created three distinct data subsets for further analyses (Fig. 1h).

***All networks dataset*** (739 networks): all quantitative networks after the above-described filtering and merging, regardless of original taxonomic scope or sampling methods. This allowed evaluation of global specialisation patterns analogous to previous syntheses not correcting for methodological or taxonomic differences.

***Full networks dataset*** (322 networks): only networks covering plant–pollinator communities without intentional taxonomic focus (i.e. not necessarily sampling all potential interactions in the community, which is usually impossible even with substantial effort). We excluded partial networks explicitly restricted to plant or pollinator groups, such as hummingbird-visited flowers or bee-plant interactions. This dataset provides more robust insights into community-level specialisation patterns by reducing bias from taxonomic sampling scope.

**Functional-group networks dataset** (926 networks, varying by functional groups; Table S3): to test within-group specialisation patterns, we used networks originally sampled with a taxonomic bias or subnetworks extracted from entire-community networks. These subnetworks were created separately for each functional pollinator group, retaining only those with ≥5 plant and ≥5 pollinator species (Table S3). Due to insufficient data, reptiles, bats, non-flying mammals, and hawkmoths were excluded from this dataset analyses (Fig. 1g).

### Specialisation metrics

To comprehensively characterise specialisation and allow reliable comparisons across networks of different size, we used two complementary size-independent metrics capturing network-level and species-level specialisation (Fig. 1i), using the *bipartite* package (Dormann *et al*. 2008).

**Network-level specialisation *H***_***2***_***’*** quantifies the deviation of observed interaction frequencies from random partner choice, reflecting overall interaction specialisation (Blüthgen *et al*. 2006). Values range from 0 (complete generalisation; interactions randomly distributed) to 1 (maximum specialisation; interactions highly restricted). This metric is standardised to account for variation in network size and species frequencies (Blüthgen *et al*. 2006).

**Community-mean species-level specialisation *d’*** represents the average specialisation of individual species within networks, allowing separation for particular functional groups. Analogous to *H*_*2*_*’*, this metric quantifies deviation of species from random interactions, ranging from 0 (generalist) to 1 (specialist; Blüthgen *et al*. 2006). Although *d’* is correlated with *H*_*2*_*’* when calculated for all species in the network (Blüthgen *et al*. 2006), it allows specialisation quantification for a part of species, such as plants, pollinators, or their functional groups. We calculated mean *d’* separately for pollinators (*d’*_*pollinators*_) and plants (*d’*_*plants*_). *d’*_*pollinators*_ was calculated across all three datasets and further partitioned by functional group. *d’*_*plants*_ was calculated only for the all networks and full networks datasets, but not for the functional-group network dataset to avoid biases introduced by restricting networks to a single pollinator group.

### Analyses of specialisation patterns

We fitted separate models for each of the three specialisation metrics (*H*_*2*_*’, d’*_*pollinators*_, and *d’*_*plants*_), using beta distribution bounded between 0 and 1. To minimise sampling bias (Blüthgen & Staab 2024), all analyses were weighted by network-specific sampling completeness. This was estimated as the proportion of observed to total interactions (completeness at order q = 0) using rarefaction and extrapolation implemented in the *Completeness*.*link* function from the *iNEXT* package (Chiu *et al*. 2023). Networks were generally well sampled, with 75% exceeding 0.75 completeness (Extended Data 3). The estimated completeness value of each network was directly used as its model weight, ensuring that each network contributed appropriately to the analyses.

To analyse patterns in interaction specialisation, we constructed eight model sets for each of the three specialisation metrics across the three datasets (with *d’*_*plants*_ not calculated for the functional-group networks dataset), employing two complementary modelling approaches. Generalised additive mixed models (GAMMs; Pedersen *et al*. 2019) were applied to *H*_*2*_*’* and *d’*_*plants*_ in the all networks and full networks datasets to capture overall patterns, as GAMMs allow flexible non-linear relationships without assuming a predefined shape. To incorporate functional group-specific responses, we fitted hierarchical generalised additive models (HGAMs) to *H*_*2*_*’* in the functional-group networks dataset and to *d’*_*pollinators*_ in all three datasets, as HGAMs extend GAMMs by modelling both global and group-specific responses to predictors. Following Pedersen et al. (2019), we constructed five HGAM types: global effect model (G: single common response across functional groups); group-specific common-shape model (S: functional groups’ responses differ in magnitude but share shape); group-specific different-shape model (I: each group has a distinct response curve); and two combined models with a global response alongside group-specific deviations: global and group-specific common shape model (GS), and global and group-specific different-shapes model (GI). All models were fitted using the *gam* function with beta family (*betar*, logit link) in the *mgcv* package (Wood 2011). Predictors were modelled with thin-plate regression splines (k = 10). For functional group-specific effects, we applied factor smooths (bs = “fs”) to obtain separate estimates for each group. Smoothing parameters were estimated by restricted maximum likelihood (REML), with a shrinkage penalty to allow splines to collapse to a straight line when the effective degrees of freedom approached one (Wood 2017). Model diagnostics were checked with the *gam*.*check* function to ensure no systematic patterns in the residuals.

To assess whether latitude predicts variation in interaction specialisation, we implemented a multi-stage modelling approach (Fig. 1j–l). First, we evaluated potential sampling biases (in the meaning of potential biases complicating interpretation of previous published results) by testing effects of three network descriptors (sampling focus, habitat openness, and geographic region), each as a fixed effect in separate models alongside latitude, for each specialisation metric and network dataset. Second, network descriptors with significant effects on latitudinal specialisation patterns were incorporated as random effects into the main latitudinal models (Fig. 1j). We then re-evaluated each random effect and sequentially removed those that did not remain significant. This resulted in the final set of eight models with latitude as the key predictor and only network descriptors that significantly influenced LSG retained as random effects (Table S4). This multi-stage approach accounts for dataset-specific variability in methodological biases while isolating the general latitudinal trend in specialisation.

To further evaluate drivers of LSG, we tested the importance of climatic, environmental, and biotic variables (Fig. 1l; Table S2). We first calculated pairwise correlations among all variables (Spearman’s ρ; Fig. S2) and removed highly correlated predictors (r > 0.7) to reduce redundancy. Because elevational variation can disrupt latitudinal patterns, we also tested the relationship of MAT to elevation, latitude, and their interaction using a linear model (*lm* function). MAT was significantly predicted by this model (p < 0.001; R^2^ = 0.89; Extended Data 1), and we therefore excluded elevation from subsequent analyses. Among the biotic variables, we excluded predicted species richness for birds, nectarivorous birds, and butterflies, as these predictors showed very limited variability within the network datasets, probably because of geographical clustering of datasets containing these functional groups, making them unsuitable for robust model testing.

We then evaluated the remaining climatic (MAT, SD_MMT, MAP), environmental (NDVI and SD_NDVI), and biotic (one predicted and eleven observed species richness measures) variables. Because biotic predictors are meaningful at different metric–dataset scales, we deployed them selectively: observed plant species richness was considered broadly for all metrics and datasets; predicted plant species richness, observed pollinator species richness and observed total species richness were considered for *H*_*2*_*’* and *d’*_*plants*_ in the all networks and full networks datasets; and observed richness of functional pollinator groups was used for species-level metrics (*d’*_*plants*_, *d’*_*pollinators*_) across datasets and additionally for *H*_*2*_*’* in the functional-group dataset (see Table S6 for details). Following the procedure for the latitudinal models (see above), we selected relevant methodological biases as random effects for each model and retained only those with significant effects. Consequently, each predictor was tested in a separate model, with MAT and MAP also examined as an interaction term. Within each of the eight model sets we compared models by AIC against the latitudinal model, treated as a null baseline. AIC values were standardised within each model set by setting the latitudinal model to zero, and we report ΔAIC values relative to this baseline (Table S6).

## Extended data

**Extended Data 1.**
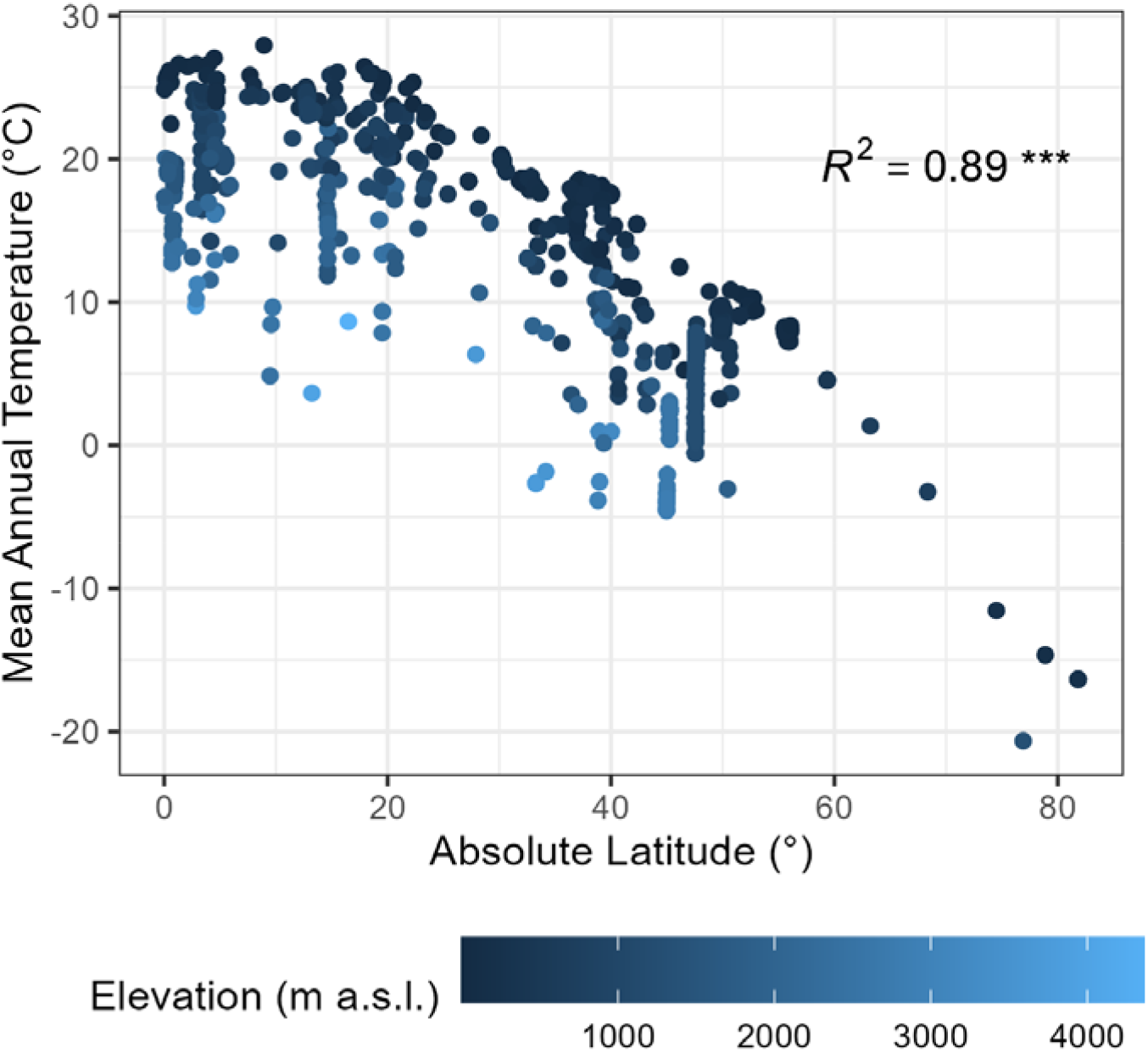
Mean annual temperature as a function of absolute latitude and elevation for the analysed plant-pollinator interaction networks. Sampling sites are coloured by their elevation. Adjusted *R*^*2*^ and p-value (***p < 0.001) originate from a linear model predicting mean annual temperature by latitude and elevation.

**Extended Data 2.**
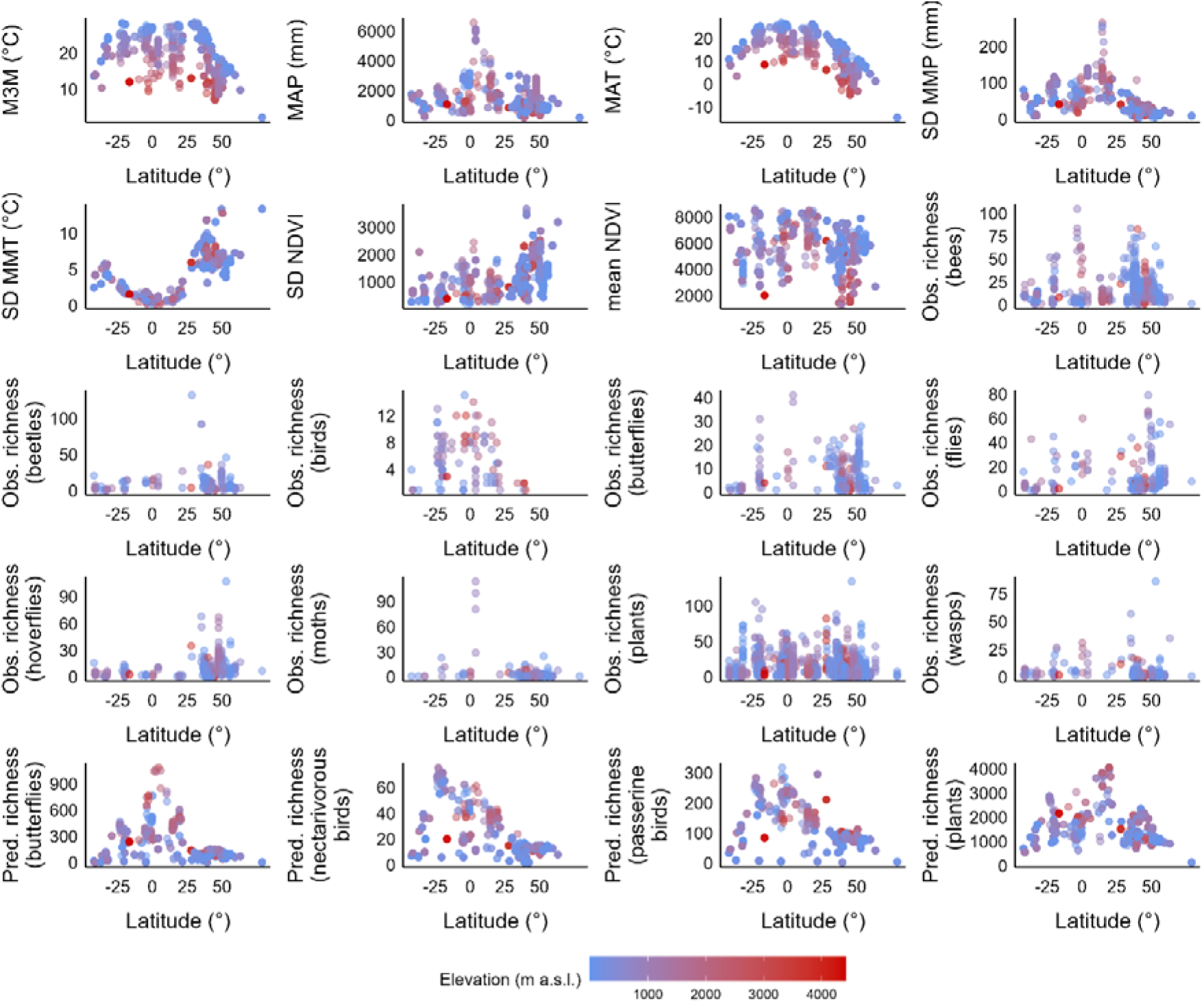
Latitudinal distribution of plant–pollinator network descriptors. Points show descriptor values for individual networks in the all networks dataset, coloured by site elevation. See Table S2 for descriptor definitions, abbreviations, and units.

**Extended Data 3.**
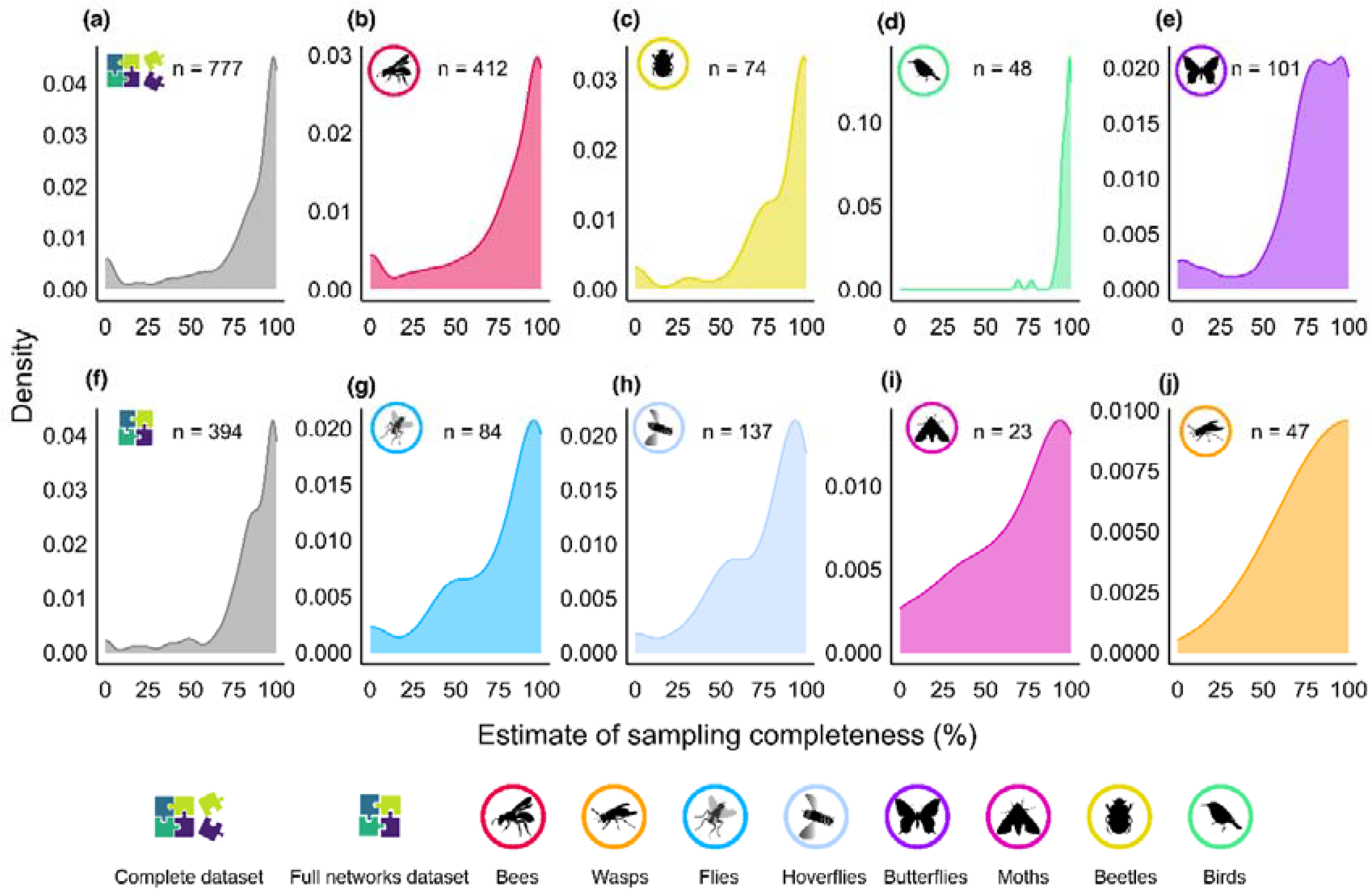
Kernel density distributions of sampling completeness (%). Panels show sampling completeness for plant–pollinator networks in the (a) all networks dataset, (f) full networks dataset, and (b–e, g–j) for individual functional groups from the functional-group networks dataset. Panel labels report the number of networks in each dataset (n).

