## Supplementary Material for "Climate-driven specialisation in plant–pollinator networks peaks outside the tropics"

Sakhalkar et al.

### List of supplementary materials:

**Figure S1.** Visual summary of latitudinal trends in specialisation stratified by methodological descriptors.
**Figure S2.** Spearman's rank correlation (ρ) matrix among plant–pollinator network descriptors.
**Table S1.** List of studies included in the dataset with their basic characteristics.
**Table S2.** Descriptors of individual plant–pollinator networks.
**Table S3.** Numbers of plant–pollinator networks by pollinator group in the functional-group networks dataset.
**Table S4.** Effects of potential methodological on latitudinal specialisation across datasets and metrics.
**Table S5.** Effects of latitude on plant–pollinator specialisation across datasets and metrics.
**Table S6.** Effects of potential methodological biases on predictor–specialisation relationships across datasets and metrics.
**Table S7.** Model comparisons identifying the best predictors of specialisation across datasets and metrics.

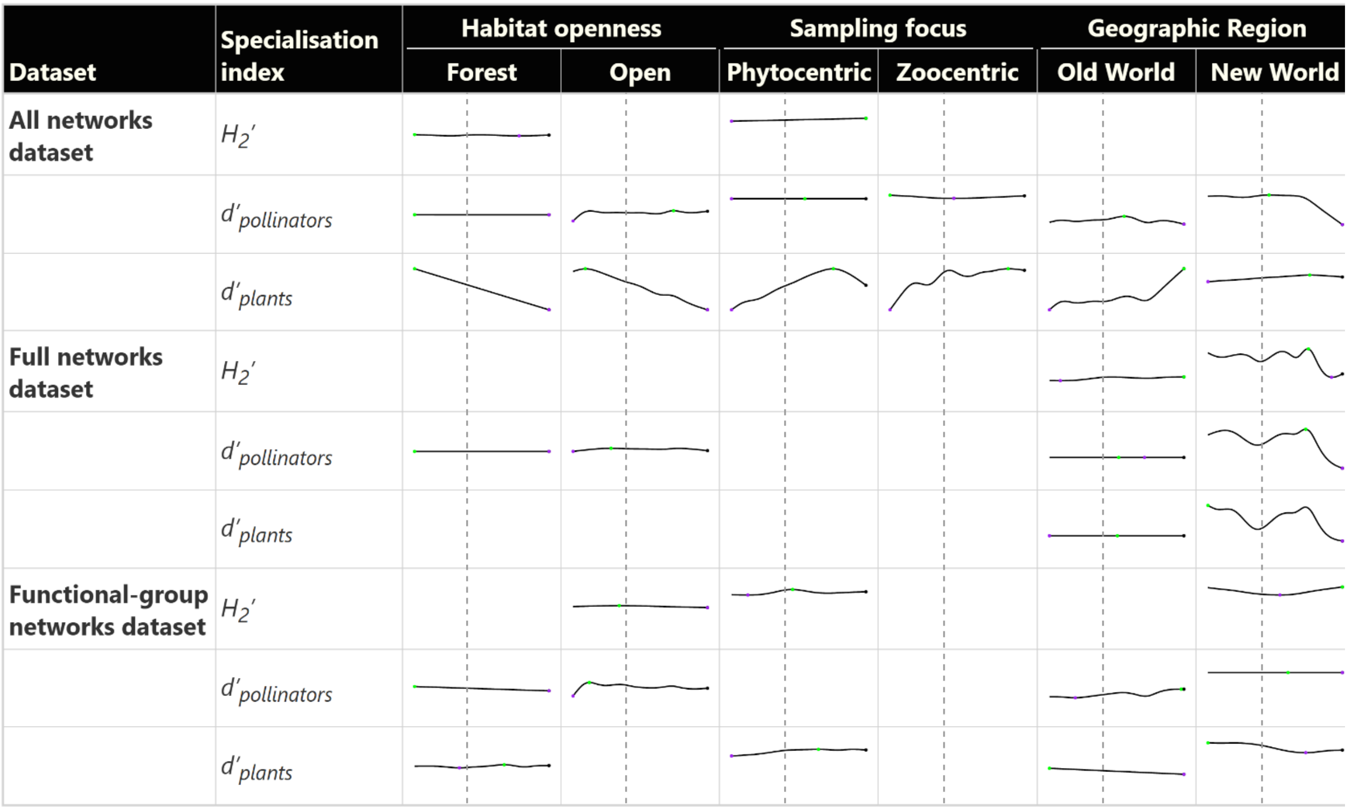


**Figure S1. Visual summary of latitudinal trends in specialisation stratified by methodological descriptors.** Mini-panels show GAMM/HGAM fitted smooths of latitude for each dataset (all, full, functional-group networks) and metric ($H_2'$, *d'pollinators*, *d'plants*) within each descriptor level. Curves are shown only when the latitude smooth is significant; otherwise, the cell is blank. The dashed vertical line marks the equator. See Online Methods for modelling details.

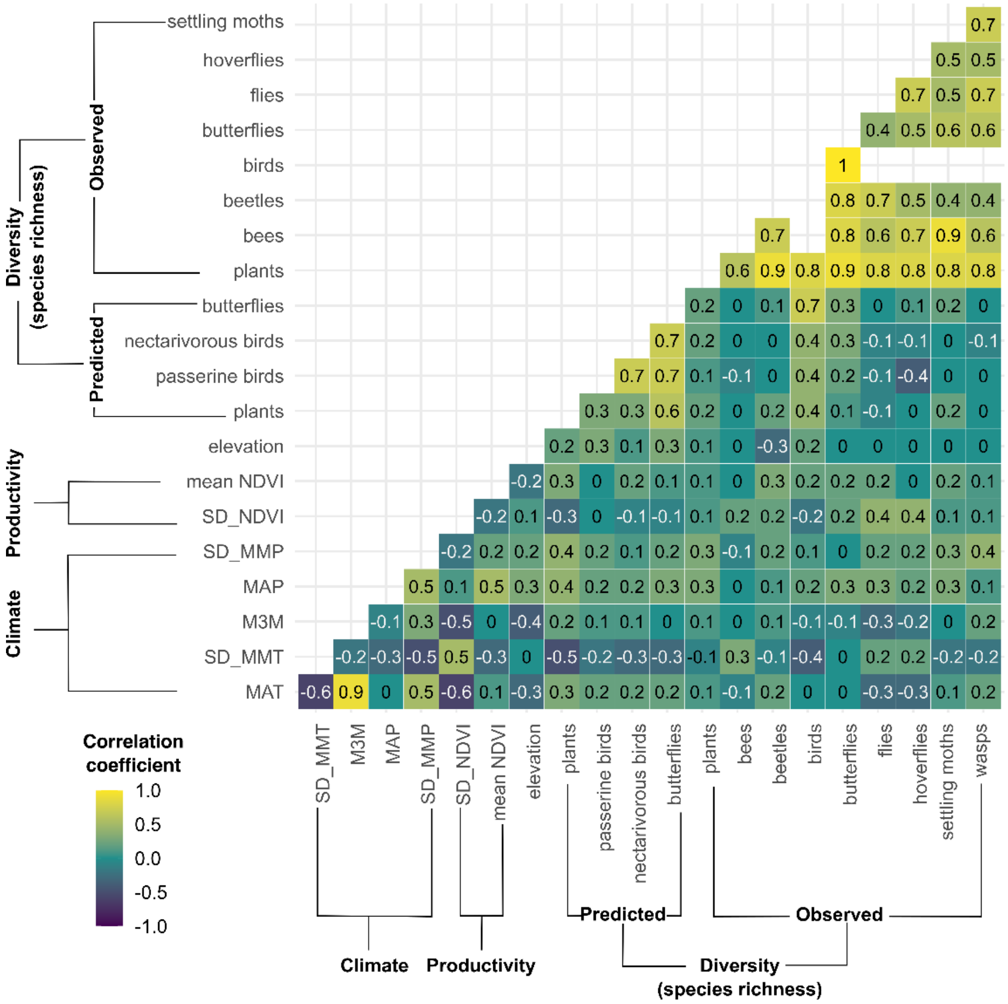


**Figure S2. Spearman's rank correlation (ρ) matrix among plant–pollinator network descriptors.** Descriptors are ordered by category (Climate, Productivity, Diversity, Elevation) as labelled. Each cell reports the pairwise **ρ**. The colour gradient indicates the direction and strength of association. Blank cells indicate insufficient overlap to estimate ρ. See Table S2 for variable definitions and abbreviations.

**Table S1. List of studies included in the dataset with their basic characteristics.** For unpublished data, their authors are listed alphabetically. For multi-network studies, geographic coordinates are the mean of network locations and elevation is reported as the range (min–max). Network completeness is defined as taxonomically restricted (partial) vs unrestricted (full). The taxonomic focus column applies only to partial networks (N/A indicates full networks).

| Dataset | Data source | Country | Geographic coordinates (°) | Elevation (m a.s.l.) | Ecoregion(s) | Habitat openness | Network completeness | Taxonomic focus (for partial networks) |
|---|---|---|---|---|---|---|---|---|
| **F_0008** | Kaiser-Bunbury et al. (2010) | Mauritius | 20.42°S, 57.44°E | 680 | Mascarene forests | open | full | N/A |
| **F_0015** | Motten (1986) | USA | 36.08°N, 79.00°W | 151 | Appalachian Piedmont forests | forest | full | N/A |
| **F_0016** | Olesen et al. (2002) | Portugal | 39.45°N, 31.19°W | 646 | Azores temperate mixed forests | open | full | N/A |
| **F_0018** | Percival (1974) | Jamaica | 17.92°N, 76.19°W | 9 | Jamaican dry forests | open | partial | flowering plants, birds, butterflies, bees |
| **F_0020** | Ponisio et al. (2017) | USA | 37.43°N, 120.37°W | 171 | California Central Valley grasslands | open | full | N/A |
| **F_0022** | Varassin & Sazima (2012) | Brazil | 19.95°S, 40.53°W | 696 | Bahia coastal forests | forest | partial | Bromeliaceae, visitors |
| **F_0025** | Dupont & Olesen (2009) | Denmark | 56.08°N, 9.18°E | 53–84 | European Atlantic mixed forests | open | full | N/A |
| **F_0027** | Inoue et al. (1990) | Japan | 35.17°N, 135.87°E | 433 | Taiheiyo evergreen forests | forest | full | N/A |
| **F_0028** | Inouye & Pyke (1988) | Australia | 36.45°S, 148.27°E | 2081 | Australian Alps montane grasslands | open | full | N/A |

| | | | | | | | | |
|---|---|---|---|---|---|---|---|---|
| **F_0029** | Kakutani et al. (1990) | Japan | 35.03°N, 135.78°E | 59 | Taiheiyo evergreen forests | forest | full | N/A |
| **F_0030** | Kato & Miura (1996) | Japan | 35.65°N, 136.08°E | 8 | Nihonkai evergreen forests | open | full | N/A |
| **F_0031** | Kato et al. (1990) | Japan | 35.33°N, 135.75°E | 818 | Nihonkai montane deciduous forests | forest | full | N/A |
| **F_0032** | Kato (2000) | Japan | 28.38°N, 129.49°E | 9 | Nansei Islands subtropical evergreen forests | open | full | N/A |
| **F_0034** | Memmott (1999) | England | 51.57°N, 2.59°W | 8 | English Lowlands beech forests | open | full | N/A |
| **F_0039** | Vázquez & Simberloff (2003) | Argentina | 41.06°S, 71.53°W | 787 | Valdivian temperate forests | forest | full | N/A |
| **F_0041** | Dicks et al. (2002) | UK | 52.59°N, 0.24°W | 2–124 | Celtic broadleaf forests | open | full | N/A |
| **F_0042** | Ollerton (2003) | South Africa | 29.17°S, 30.13°E | 1392 | Drakensberg grasslands | open | partial | Asclepiadaceae, visitors |
| **F_0046** | Small (1976) | Canada | 45.40°N, 75.5°W | 68 | Eastern Great Lakes lowland forests | open | full | N/A |
| **F_0047** | Smith-Ramirez et al. (2005) | Chile | 42.00°S, 73.58°W | 80 | Valdivian temperate forests | forest | full | N/A |
| **F_0049** | Vazquez (2002) | Argentina | 41.08°S, 71.53°W | 829 | Valdivian temperate forests | forest | full | N/A |
| **F_0050** | Vizentin-Bugoni et al. (2015) | Brazil | 23.34°S, 45.12°W | 966 | Serra do Mar coastal forests | forest | partial | ornithophilous plants, hummingbirds |
| **F_0051** | Abreu & Vieira (2004) | Brazil | 20.75°S, 42.92°W | 709 | Bahia interior forests | forest | partial | ornithophilous plants, hummingbirds |

| | | | | | | | | |
|---|---|---|---|---|---|---|---|---|
| **F_0052** | del Coro Arizmendi & Ornelas (1990) | Mexico | 19.50°N, 105.05°W | 86 | Jalisco dry forests | forest | partial | ornithophilous plants, hummingbirds |
| **F_0053** | Canela (2006) | Brazil | 22.50°S, 44.83°W | 793 | Alto Paraná Atlantic forests | forest | partial | ornithophilous plants, hummingbirds |
| **F_0054** | Las-Casas et al. (2012) | Brazil | 7.87°S, 36.40°W | 563 | Caatinga | open | partial | ornithophilous plants, hummingbirds |
| **F_0055** | Aquiles Gutierrez et al. (2004) | Colombia | 1.25°N, 77.43°W | 2709 | Northwest Andean montane forests | forest | partial | ornithophilous plants, hummingbirds |
| **F_0056** | Lara (2006) | Mexico | 19.23°N, 98.97°W | 2239 | Central Mexican matorral | forest | partial | ornithophilous plants, hummingbirds |
| **F_0057** | Lasprilla & Sazima (2004) | Colombia | 0.04°N, 72.27°W | 160 | Caqueta moist forests | forest | partial | ornithophilous plants, hummingbirds |
| **F_0060** | Martínez-Núñez & Rey (2020) | Spain | 37.42°N, 4.56°W | 428–1313 | Iberian sclerophyllous and semi-deciduous forests | open | partial | flowering plants, bees |
| **F_0061** | Unpublished: Š. Janeček, I.N. Kobe, S.P. Sakhalkar, A. Sounapoglou, R. Tropek | Czechia | 50.65°N, 15.78°E | 513–1075 | Western European broadleaf forests | forest | full | N/A |
| **F_0062** | Klomberg et al. (2022) | Cameroon | 4.12°N, 9.09°E | 650–2200 | Mount Cameroon and Bioko montane forests | forest | full | N/A |
| **F_0063** | Unpublished: M. Bartoš, Š. Janeček, | Czechia | 49.78°N, 15.82°E | 450–1000 | Western European | open | full | N/A |

| | | | | | | | | |
|---|---|---|---|---|---|---|---|---|
| | J. Jersáková, R. Tropek | | | | broadleaf forests; Central European mixed forests | | | |
| **F_0064** | Hernández-Castellano et al. (2021) | Spain | 41.73°N, 2.41°E | 418 | Northeast Spain and Southern France Mediterranean forests | forest | full | N/A |
| **F_0065** | Reverté et al. (2019) | Spain | 41.28°N, 1.89°E | 402 | Northeast Spain and Southern France Mediterranean forests | open | full | N/A |
| **F_0067** | Unpublished: T.C. Ling | China | 21.62°N, 101.58°E | 835 | Northern Indochina subtropical forests | forest | full | N/A |
| **F_0068** | Unpublished: W. Dáttilo, P. Luna | Mexico | 19.46°N, 96.85°W | 14–3275 | Mesoamerican Gulf-Caribbean mangroves; Trans-Mexican Volcanic Belt pine-oak forests; Veracruz dry forests; Veracruz moist forests; Oaxacan montane forests | forest | partial | flowering plants, bees |
| **F_0069** | Santamaría Bueno & Méndez Iglesias (2021) | Spain | 41.17°N, 4.37°W | 2145–2844 | Cantabrian mixed forests; Iberian conifer forests | open | full | N/A |
| **F_0070** | Janeček et al. (2021) | Cameroon | 4.12°N, 9.09°E | 650–2200 | Mount Cameroon and Bioko montane forests | forest | partial | flowering plants, nectarivorous birds |

| | | | | | | | | |
|---|---|---|---|---|---|---|---|---|
| **F_0071** | Rech & Lúcia Absy (2011) | Brazil | 0.69°S, 63.37°W | 6–87 | Negro-Branco moist forests; Monte Alegre várzea; Rio Negro campinarana; Japurá-Solimões-Negro moist forests; Caatinga | forest | partial | flowering plants, nectarivorous birds |
| **F_0072** | Unpublished: J.P.R. Borges, A.C.P. Machado | Brazil | 18.2°S, 43.57°W | 1354 | Cerrado | open | partial | Asteraceae, nectarivorous birds |
| **F_0073** | Unpublished: M.S. Amorim, A.R. Rech | Brazil | 18.17°S, 43.46°W | 994–1354 | Cerrado | open | partial | flowering plants, hummingbirds |
| **F_0074** | Unpublished: F.D. Reis | Brazil | 14.85°S, 43.94°W | 453 | Caatinga | forest | partial | flowering plants, bats |
| **F_0075** | Nielsen & Totland (2014) | Norway | 59.35°N, 9.75°E | 488 | Scandinavian and Russian taiga | forest | full | N/A |
| **F_0076** | Matsubara et al. (2023) | Japan | 40.67°N, 140.87°E | 572–1304 | Honshu alpine conifer forests; Nihonkai montane deciduous forests | open | full | N/A |
| **F_0077** | Unpublished: C.S. Ballarin | Brazil | 22.94°S, 48.46°W | 854 | Cerrado | open | partial | flowering plants, bees |
| **F_0082** | Dzekashu et al. (2023) | Kenya | 2.05°S, 37.65°E | 522–2515 | East African montane forests; Northern Acacia-Commiphora bushlands and thickets; Eastern Arc forests | forest | full | N/A |

| | | | | | | | | |
|---|---|---|---|---|---|---|---|---|
| **F_0083** | Crowther et al. (2014) | UK | 52.53°N, 1.30°E | 9–66 | Celtic broadleaf forests; English Lowlands beech forests | open | partial | flowering plants, bees |
| **F_0085** | Unpublished: J.M. Cook, A.M. Gilpin | Australia | 33.42°S, 150.01°E | 450–880 | Southeast Australia temperate forests; Eastern Australian temperate forests | open | full | N/A |
| **F_0086** | Hopfenmüller et al. (2014) | Germany | 49.92°N, 11.30°E | 348–533 | Western European broadleaf forests | open | partial | flowering plants, wild bees |
| **F_0087** | Bezerra et al. (2009) | Brazil | 8.71°S, 37.20°W | 625 | Cerrado | forest | partial | Malpighiaceae, wild bees |
| **F_0088** | Diniz & Aguiar (2023) | Brazil | 15.68°S, 47.98°W | 1090 | Cerrado | open | partial | flowering plants, bats |
| **F_0092** | Arroyo-Correa et al. (2023) | Spain | 37.01°N, 6.51°W | 22–26 | Southwest Iberian Mediterranean sclerophyllous and mixed forests | open | full | N/A |
| **F_0093** | Simanonok & Burkle (2014) | USA | 44.99°N, 109.42°W | 2863–3287 | South Central Rockies forests | open | full | N/A |
| **F_0094** | Figueroa et al. (2020) | USA | 42.66°N, 76.69°W | 114 | Eastern Great Lakes lowland forests | open | partial | flowering plants, Bombus |
| **F_0095** | Unpublished: C. Bosenbecker, P.K. Maruyama | Brazil | 19.86°S, 44.01°W | 813–1240 | Bahia interior forests; Cerrado; Campos Rupestres montane savanna | forest | partial | flowering plants, Bombus |

| | | | | | | | | |
|---|---|---|---|---|---|---|---|---|
| **F_0096** | Unpublished: I.N. Gomes, P.K. Maruyama, V.H.D. Silva | Brazil | 19.91°S, 43.95°W | 813–1240 | Bahia interior forests; Cerrado; Campos Rupestres montane savanna | open | partial | flowering plants, Bombus |
| **F_0097** | Ribas-Marqués et al. (2022) | Spain | 39.62°N, 2.98°E | 164 | Northeast Spain and Southern France Mediterranean forests | forest | partial | flowering plants, Bombus |
| **F_0098** | Unpublished: R.N. Afagwu, A.M.T. Castagna, M.M. Mayberry, A.L. Russell | USA | 37.47°N, 93.9°W | 264–359 | Ozark Highlands mixed forests; Central Tallgrass prairie | open | full | N/A |
| **F_0099** | Primack (1983) | New Zealand | 43.15°S, 171.48°E | 600–1800 | New Zealand South Island montane grasslands | open | full | N/A |
| **F_0103** | Magrach et al. (2023) | Spain | 40.15°N, 4.56°W | 31–1097 | Cantabrian mixed forests; Southwest Iberian Mediterranean sclerophyllous and mixed forests | open | full | N/A |
| **F_0105** | CaraDonna (2020) | USA | 38.96°N, 106.99°W | 2898 | Colorado Rockies forests | forest | partial | flowering plants, insects, hummingbirds |
| **F_0107** | Hervías-Parejo et al. (2023) | Spain | 39.93°N, 4.17°E | 23–53 | Northeast Spain and Southern France Mediterranean forests | forest | full | N/A |

| | | | | | | | | |
|---|---|---|---|---|---|---|---|---|
| **F_0108** | Junker et al. (2010) | Germany | 49.79°N, 9.94°E | 188 | Western European broadleaf forests | open | full | N/A |
| **F_0109** | Bain et al. (2022) | China | 38.96°N, 106.99°E | 1234 | Ordos Plateau steppe | open | full | N/A |
| **F_0110** | Unpublished: J. Ollerton, S. Tarrant | England | 52.27°N, 0.88°W | 124 | Celtic broadleaf forests | open | full | N/A |
| **F_0111** | Ornai & Keasar (2020) | Israel | 32.74°N, 35.04°E | 494 | Eastern Mediterranean conifer-broadleaf forests | forest | partial | flowering plants, visitors |
| **F_0112** | Prendergast (2020) | Australia | 31.99°S, 115.85°E | 5–42 | Southwest Australia woodlands | open | partial | flowering plants, visitors |
| **F_0114** | Burkle et al. (2013) | USA | 39.25°N, 89.85°W | 164–203 | Central US forest-grasslands transition | forest | partial | flowering plants, bees |
| **F_0115** | Burkle & Knight (2012) | USA | 38.37°N, 90.82°W | 146–254 | Ozark Highlands mixed forests | open | full | N/A |
| **F_0116** | Heil & Burkle (2018) | USA | 45.25°N, 110.52°W | 2013–2435 | South Central Rockies forests | open | partial | flowering plants, bees |
| **F_0117** | da Rocha-Filho et al. (2021) | Brazil | 21.17°S, 47.8°W | 546 | Alto Paraná Atlantic forests/Cerrado | forest | partial | flowering plants, bees |
| **F_0118** | Seitz et al. (2020) | USA | 39.02°N, 76.83°W | 40–63 | Southeast US conifer savannas | open | partial | flowering plants, bees |
| **F_0121** | Abreu & Vieira (2004) | Brazil | 20.75°S, 42.92°W | 709 | Bahia interior forests | forest | partial | ornithophilous plants, bees |
| **F_0124** | Araújo et al. (2013) | Brazil | 19.95°S, 40.52°W | 671 | Bahia coastal forests | forest | partial | ornithophilous plants, bees |
| **F_0125** | del Coro Arizmendi & Ornelas (1990) | Mexico | 19.50°N, 105.05°W | 86 | Jalisco dry forests | forest | partial | ornithophilous plants, bees |

| | | | | | | | | |
|---|---|---|---|---|---|---|---|---|
| **F_0126** | Barbosa-Filho & Araujo (2013) | Brazil | 20.51°S, 54.62°W | 549 | Cerrado | forest | partial | flowering plants, bees |
| **F_0127** | Buzato et al. (2000) | Brazil | 23.38°S, 45.12°W | 6–1257 | Serra do Mar coastal forests | forest | partial | ornithophilous plants, bees |
| **F_0128** | Canela (2006) | Brazil | 22.5°S, 44.83°W | 793 | Alto Paraná Atlantic forests | forest | partial | ornithophilous plants, bees |
| **F_0129** | Cotton(1998) | Colombia | 3.82°S, 70.27°W | 69 | Iquitos várzea | forest | partial | flowering plants, bees |
| **F_0133** | Gonzalez & Loiselle (2016) | Peru | 9.71°S, 76.16°W | 3287 | Peruvian Yungas | open | partial | all flowering plants, plus focus on ornithophilous plants, bees |
| **F_0135** | Aquiles Gutierrez et al. (2004) | Colombia | 1.25°N, 77.43°W | 2709 | Northwest Andean montane forests | forest | partial | ornithophilous plants, bees |
| **F_0136** | Las-Casas et al. (2012) | Brazil | 7.87°S, 36.4°W | 563 | Caatinga | open | partial | ornithophilous plants, bees |
| **F_0137** | Lasprilla & Sazima (2004) | Colombia | 0.07°N, 72.45°W | 131 | Caqueta moist forests | forest | partial | flowering plants, bees |
| **F_0140** | Machado (2009) | Brazil | 13.12°S, 41.58°W | 1177 | Caatinga | open | partial | flowering plants, bees |
| **F_0141** | Machado (2014) | Brazil | 13.12°S, 41.57°W | 1143 | Caatinga | open | partial | ornithophilous plants, bees |
| **F_0142** | Maglianesi et al. (2014) | Costa Rica | 10.30°N, 84.07°W | 57–1987 | Isthmian-Atlantic moist forests; Talamancan montane forests | forest | partial | ornithophilous plants, bees |
| **F_0143** | Martínez-García & Ortiz-Pulido (2014) | Mexico | 20.61°N, 98.75°W | 1676–2307 | Sierra Madre Oriental pine-oak forests | open | partial | flowering plants, bees |

| | | | | | | | | |
|---|---|---|---|---|---|---|---|---|
| **F_0144** | Maruyama et al. (2015) | Brazil | 23.34°S, 44.87°W | 132–149 | Serra do Mar coastal forests | forest | partial | flowering plants, bees |
| **F_0147** | Partida-Lara et al. (2018) | Mexico | 15.63°N, 92.82°W | 645–2091 | Sierra Madre de Chiapas moist forests | forest | partial | flowering plants, bees |
| **F_0148** | Ramírez-Burbano et al. (2017) | Colombia | 2.59°N, 76.97°W | 1747–2460 | Northwest Andean montane forests | forest | partial | flowering plants, bees |
| **F_0149** | Rodrigues & Araujo (2011) | Brazil | 19.84°S, 49.08°W | 603–1285 | Cerrado; Campos Rupestres montane savanna | forest | partial | flowering plants, bees |
| **F_0150** | Sazima et al. (1996) | Brazil | 22.73°S, 45.58°W | 1616 | Serra do Mar coastal forests | open | partial | flowering plants, bees |
| **F_0151** | Snow & Snow (1972) | Trinidad & Tobago | 10.67°N, 61.28°W | 322 | Trinidad and Tobago moist forest | forest | partial | flowering plants, bees |
| **F_0152** | Snow & Snow (1979) | Colombia | 5.45°N, 73.6°W | 1689–2617 | Magdalena Valley montane forests; Cordillera Oriental montane forests | open | partial | flowering plants, bees |
| **F_0153** | Snow & Snow (1986) | Brazil | 23.63°S, 45.85°W | 951 | Serra do Mar coastal forests | open | partial | flowering plants, bees |
| **F_0154** | Sonne et al. (2019) | Ecuador | 4.06°S, 79.07°W | 1300–2951 | Eastern Cordillera Real montane forests | forest | partial | flowering plants with a minimal corolla tube, bees |
| **F_0155** | Tinoco et al. (2016) | Ecuador | 2.89°S, 79.12°W | 3029–3347 | Eastern Cordillera Real montane forests | forest | partial | all but herbaceous Asteraceae, bees |

| | | | | | | | | |
|---|---|---|---|---|---|---|---|---|
| **F_0157** | Vizentin-Bugoni et al. (2015) | Brazil | 23.28°S, 45.05°W | 1077 | Serra do Mar coastal forests | forest | partial | flowering plants, bees |
| **F_0161** | de Araújo et al. (2011) | Brazil | 18.99°S, 48.3°W | 840 | Cerrado | open | partial | flowering plants, bees |
| **F_0162** | Bartomeus et al. (2008) | Spain | 42.32°N, 3.30°E | 74 | Northeast Spain and Southern France Mediterranean forests | open | full | N/A |
| **F_0163** | Shay et al. (2016) | USA | 21.55°N, 158.2°W | 398 | Hawai'i tropical moist forests | open | full | N/A |
| **F_0164** | Benadi et al. (2013) | Germany | 47.65°N, 13.01°E | 522–1441 | Alps conifer and mixed forests | open | partial | flowering plants, bees |
| **F_0167** | Probert (2019) | New Zealand | 36.96°S, 174.47°E | 58–179 | Northland temperate kauri forests | forest | partial | flowering plants, bees |
| **F_0168** | Peralta et al. (2020) | Argentina | 32.53°S, 68.95°W | 1239 | High Monte | open | full | N/A |
| **F_0169** | Robinson et al. (2018) | Canada | 78.88°N, 75.8°W | 1 | Canadian High Arctic tundra | open | full | N/A |
| **F_0170** | Robson (2014) | Canada | 49.71°N, 99.10°W | 345 | Canadian Aspen forests and parklands | open | full | N/A |
| **F_0171** | Mendonça Santos et al. (2010) | Brazil | 12.7°S, 39.77°W | 262 | Caatinga | open | partial | flowering plants, bees |
| **F_0172** | Song (2015) | Mongolia | 50.44°N, 100.39°E | 1715 | Sayan montane conifer forests | open | full | N/A |
| **F_0173** | Souza et al. (2018) | Brazil | 20.03°S, 55.34°W | 80–646 | Cerrado; Humid Chaco; Pantanal | open | full | N/A |
| **F_0174** | Fang & Huang (2016) | China | 27.90°N, 99.64°E | 3317 | Hengduan Mountains | open | full | N/A |

| | | | | | | | | |
|---|---|---|---|---|---|---|---|---|
| | | | | | subalpine conifer forests | | | |
| **F_0175** | Lara-Romero et al. (2016) | Spain | 40.83°N, 3.95°W | 1766 | Iberian conifer forests | open | full | N/A |
| **F_0176** | Guy et al. (2021) | Kenya | 0.38°N, 36.88°E | 1591–1733 | Northern Acacia-Commiphora bushlands and thickets | open | full | N/A |
| **F_0178** | Resasco et al. (2021) | USA | 40.03°N, 105.54°W | 2930 | Colorado Rockies forests | open | full | N/A |
| **F_0179** | Escobedo-Kenefic et al. (2020) | Guatemala | 14.58°N, 90.76°W | 748–1777 | Central American pine-oak forests; Central American montane forests | forest | partial | flowering plants, bees |
| **F_0180** | Kaiser-Bunbury et al. (2017) | Seychelles | 4.67°S, 55.47°E | 331–448 | Granitic Seychelles forests | forest | full | N/A |
| **F_0182** | Ballantyne et al. (2017) | Israel | 32.73°N, 35.01°E | 167 | Eastern Mediterranean conifer-broadleaf forests | open | full | N/A |
| **F_0183** | Ballantyne et al. (2015) | UK | 50.71°N, 2.21°W | 59 | English Lowlands beech forests | open | full | N/A |
| **F_0184** | Unpublished: G. Nakas, T. Petanidou | Greece | 38.24°N, 26.02°E | 51–200 | Aegean and Western Turkey sclerophyllous and mixed forests | open | full | N/A |
| **F_0185** | Unpublished: T. Petanidou | Greece | 37.81°N, 25.29°E | 36–375 | Aegean and Western Turkey sclerophyllous and mixed forests | open | full | N/A |
| **F_0186** | Librán-Embid et al. (2021) | Germany | 51.55°N, 9.95°E | 171 | Western European broadleaf forests | open | full | N/A |

| | | | | | | | | |
|---|---|---|---|---|---|---|---|---|
| **F_0187** | Unpublished: A.D. Bjorkman, W.H.A. Osterman | Sweden | 63.21°N, 12.39°E | 723 | Scandinavian Montane Birch forest and grasslands | open | full | N/A |
| **F_0188** | Balmaki et al. (2022) | USA | 39.43°N, 119.58°W | 1376–3151 | Sierra Nevada forests; Great Basin shrub steppe | open | partial | flowering plants, bees, flies |
| **F_0189** | Unpublished: V. De Cardenas, S. Lozada-Gobilard | Bolivia | 16.49°S, 68.12°W | 3674 | Central Andean wet puna | forest | full | N/A |
| **F_0190** | Unpublished: Y. Clough, V. Hederström, T. Krausl | Sweden | 55.73°N, 13.71°E | 34–124 | Baltic mixed forests | open | full | N/A |
| **F_0191** | Querejeta et al. (2023) | France | 46.15°N, 0.42°W | 75 | European Atlantic mixed forests | open | partial | flowering plants, bees, flies |
| **F_0192** | Maihoff et al. (2023) | Germany | 47.58°N, 12.90°E | 646–2005 | Alps conifer and mixed forests | open | partial | flowering plants, bees, flies |
| **F_0193** | van der Kooi et al. (2016) | Netherlands | 53.00°N, 6.58°E | 12 | European Atlantic mixed forests | open | full | N/A |
| **F_0194** | Marcacci et al. (2023) | India | 13.00°N, 77.56°E | 696–933 | Deccan thorn scrub forests; South Deccan Plateau dry deciduous forests | open | partial | flowering plants, bees, flies |
| **F_0195** | Unpublished: K.T. Burghardt, K.R. Urban-Mead | United States | 41.42°N, 72.8°W | 58 | Northeast US Coastal forests | open | full | N/A |

| | | | | | | | | |
|---|---|---|---|---|---|---|---|---|
| **F_0196** | Cortina et al. (2022) | United States | 31.72°N, 97.23°W | 171–375 | Texas blackland prairies; Cross-Timbers savanna-woodland; Edwards Plateau savanna | open | full | N/A |
| **F_0197** | Biella et al. (2017) | Italy | 44.68°N, 9.26°E | 1476–1570 | Appenine deciduous montane forests | open | full | N/A |
| **F_0199** | Escobedo-Kenefic et al. (2020) | Guatemala | 14.64°N, 90.88°W | 1666–2630 | Central American pine-oak forests | forest | partial | flowering plants, bees, flies |

**Table S2. Descriptors of individual plant–pollinator networks**, with their category, definition, units or categories, and data source/reference.

| Category | Descriptor (abbreviation) | Definition | Additional comment, units, or categories | Source / Reference |
|---|---|---|---|---|
| **Study and methodological details** | Data source | | Publication, database, or authors of unpublished data | Original study |
| **Study and methodological details** | Reasons for a network split in original study | Rationale for splitting datasets (distinct sites, elevations, habitats, or vertical strata) | This information was used to decide whether and how networks could be pooled for our analyses. | Original study |
| **Study and methodological details** | Field sampling method | Sampling design and main field methods used for data collection | This information was used to classify networks by type (qualitative vs. quantitative) and sampling focus (zoo- vs. phytocentric), assess taxonomic completeness (full vs. partial), and determine constraints on observed interactions at both the lower (plants) and upper (pollinators) trophic levels. | Original study |
| **Study and methodological details** | Sampling focus | | Zoocentric, phytocentric | Original study |
| **Study and methodological details** | Network completeness | | Full network, partial network | Original study |
| **Study and methodological details** | Lower-level (plant) taxonomic constraints | Specified if plants were sampled with taxonomic bias | Specified taxa or “community-wide” | Original study |
| **Study and methodological details** | Upper-level (pollinator) taxonomic constraints | Specified if pollinators were sampled with taxonomic bias | Specified taxa or “community-wide” | Original study |

| | | | | |
|---|---|---|---|---|
| **Geographic information** | Country | | | Original study |
| **Geographic information** | Latitude | Geographic coordinates | Decimal degrees | Original study |
| **Geographic information** | Longitude | Geographic coordinates | Decimal degrees | Original study |
| **Geographic information** | Absolute latitude | Absolute value of latitude | Decimal degrees | Original study |
| **Geographic information** | Elevation | | Metres above sea level | Original study |
| **Geographic information** | Region | | New World, Old World | Original study |
| **Environmental conditions** | Habitat openness | Predominant habitat openness at sampling site | Forest, open habitat | Original study |
| **Environmental conditions** | Mean annual temperature (MAT) | | °C | Fick and Hijmans (2017) |
| **Environmental conditions** | Standard deviation of mean monthly temperature (SD_MMT) | | ±°C | Fick and Hijmans (2017) |
| **Environmental conditions** | Mean temperature of the three warmest months (M3M) | | °C | Fick and Hijmans (2017) |
| **Environmental conditions** | Mean annual precipitation (MAP) | | mm | Fick and Hijmans (2017) |
| **Environmental conditions** | Standard deviation of mean monthly precipitation (SD_MMP) | | ±mm | Fick and Hijmans (2017) |
| **Environmental conditions** | Mean annual Normalised Difference Vegetation Index (mean NDVI) | | NDVI index values | Didan (2015) |
| **Environmental conditions** | Standard deviation of mean monthly Normalised Difference Vegetation Index (SD_NDVI) | | NDVI index values | Didan (2015) |
| **Diversity** | Predicted plant species richness | | Number of species | Cai et al. (2023) |

| | | | | |
|---|---|---|---|---|
| **Diversity** | Predicted bird species richness | | Number of species | Jenkins et al. (2013) |
| **Diversity** | Predicted nectarivorous bird species richness | | Number of species | Dalsgaard & Ollerton (unpublished species list used to select rangemaps from BirdLife data) |
| **Diversity** | Predicted butterfly species richness | | Number of species | Daru (2024) |
| **Diversity** | Observed plant species richness | As recorded in the studied network | Number of species | Original study |
| **Diversity** | Observed pollinator species richness | As recorded in the studied network | Number of species | Original study |
| **Diversity** | Observed total species richness | As recorded in the studied network | Number of species | Original study |
| **Diversity** | 8 predictors of Observed species richness of functional groups | As recorded in the studied network | Number of species | Original study |

**Table S3. Numbers of plant–pollinator networks by pollinator group in the functional-group networks dataset.**

| Functional group | Number of networks |
|---|---|
| **Bees** | 412 |
| **Beetles** | 74 |
| **Birds** | 48 |
| **Butterflies** | 101 |
| **Flies** | 84 |
| **Hoverflies** | 137 |
| **settling moths** | 23 |
| **wasps** | 47 |

**Table S4**. **Effects of potential methodological on latitudinal specialisation across datasets and metrics.** Results are shown for network-level $H_2'$ and community-mean species-level $d'_{plants}$ and $d'_{pollinators}$. For each descriptor (habitat openness, region, sampling focus), HGAMs were fitted with latitude modelled within descriptor levels; cells report the estimated degrees of freedom (edf) for the latitude smooth. edf = 0 indicates no effect (smooth shrunk to zero), edf ≈ 1 an approximately linear effect, and edf > 1 a non-linear effect (see Fig. S1 for individual smooth curves). Significant descriptor effects were used as random effects in the final latitudinal models (see Online Methods). Significance refers to the latitude smooth term: * $p < 0.05$, ** $p < 0.01$, *** $p < 0.001$. See Table S2 for descriptor definitions.

| Dataset | Specialization metric | Habitat | | Region | | Sampling focus | |
|---|---|---|---|---|---|---|---|
| | | Forest | Open | New World | Old World | Phytocentric | Zoocentric |
| All networks | $H_2'$ | 4.29 *** | 0 | 0.94 | 0 | 0.94 *** | 0.26 |
| | $d'_{pollinators}$ | 0 * | 8.30 *** | 6.02 *** | 7.12 *** | 0 *** | 2.21 *** |
| | $d'_{plants}$ | 0.87 *** | 6.76 *** | 0.48 *** | 8.12 *** | 4.28 *** | 6.37 *** |
| Full networks | $H_2'$ | 0 | 2.08 *** | 3.21 *** | 0 | 5.48 *** | 0 |
| | $d'_{pollinators}$ | 0.64 *** | 8.62 *** | 0 *** | 6.85 *** | 0 | 0 |
| | $d'_{plants}$ | 6.92 *** | 0 | 4.21 *** | 0.95 *** | 4.93 *** | 0 |
| Functional-group networks | $H_2'$ | 0 | 0.98 | 6.54 *** | 3.98 *** | 0 | 0 |
| | $d'_{pollinators}$ | 0 *** | 4.42 *** | 6.39 *** | 0.01 *** | 0 | 0 |

**Table S5. Effects of latitude on plant–pollinator specialisation across datasets and metrics.** Results are shown for network-level *H₂'* and community-mean species-level $d'_{plants}$ and $d'_{pollinators}$ across the all networks, full networks, and functional-group networks datasets ($d'_{plants}$ not fitted for the functional-group dataset). Cells report the estimated degrees of freedom (edf) for the latitude smooth in the final models, conditional on retained random effects; for $d'_{pollinators}$ HGAMs, group-specific edf and p-values are additionally shown for each functional group. edf = 0 indicates no effect (smooth shrunk to zero), edf ≈ 1 an approximately linear effect, and edf > 1 a non-linear effect. Models incorporated significant potential methodological biases (habitat openness, sampling focus, region) as random effects (Table S4). $R^2adj$ gives the variance explained by the whole model. P-values refer to the approximate significance of the latitude smooth: * p < 0.05, ** p < 0.01, *** p < 0.001.

| Specialisation metric | Random effects | $R^2_{adj}$ | Functional group | EDF | p-value |
|---|---|---|---|---|---|
| **All networks dataset** | | | | | |
| $H_2'$ | Region | 0.042 | | 4.76 | 0.001*** |
| $d'_{pollinators}$ | Habitat openness, region | 0.127 | | 7.52 | 0*** |
| | | | bees | 8.06 | 0*** |
| | | | beetles | 0.65 | 0.08 |
| | | | birds | 2.39 | 0*** |
| | | | butterflies | 0.54 | 0.13 |
| | | | flies | 0.00 | 0.36 |
| | | | hoverflies | 0.00 | 0.43 |
| | | | settling moths | 1.26 | 0.01** |
| | | | wasps | 0.00 | 0.621 |
| $d'_{plants}$ | none | 0.23 | | 6.05 | 0*** |
| **Full networks dataset** | | | | | |
| $H_2'$ | none | 0.002 | | 0.00 | 0.381 |
| $d'_{pollinators}$ | Region | 0.09 | | 7.09 | 0*** |
| | | | bees | 0.00 | 0.68 |
| | | | beetles | 0.70 | 0.065 |
| | | | birds | 0.65 | 0.087 |
| | | | butterflies | 0.95 | 0.02* |
| | | | flies | 0.00 | 0.608 |
| | | | hoverflies | 0.00 | 0.646 |
| | | | settling moths | 4.37 | 0*** |
| | | | wasps | 0.00 | 0.65 |
| $d'_{plants}$ | none | 0.01 | | 0.85 | 0*** |
| **Functional-group networks dataset** | | | | | |
| $H_2'$ | none | 0.220 | | 5.89 | 0*** |
| | | | bees | 2.41 | 0*** |
| | | | beetles | 5.88 | 0.00 |

| Specialisation metric | Random effects | $R^2_{adj}$ | Functional group | EDF | p-value |
|---|---|---|---|---|---|
| | | | birds | 2.44 | 0*** |
| | | | butterflies | 1.56 | 0*** |
| | | | flies | 0.00 | 0.889 |
| | | | hoverflies | 3.40 | 0.001** |
| | | | settling moths | 2.96 | 0.002** |
| | | | wasps | 2.68 | 0.663 |
| $d'_{pollinators}$ | Habitat openness | 0.33 | | 5.06 | 0*** |
| | | | bees | 8.03 | 0*** |
| | | | beetles | 0.00 | 0.73 |
| | | | birds | 2.33 | 0*** |
| | | | butterflies | 0.802 | 0.02* |
| | | | flies | 0.00 | 0.99 |
| | | | hoverflies | 7.48 | 0*** |
| | | | settling moths | 7.18 | 0*** |
| | | | wasps | Not enough data | |

**Table S6. Effects of potential methodological biases on predictor–specialisation relationships across datasets and metrics.** For each dataset (all networks, full networks, functional-group networks) and specialisation metric (network-level $H_2'$; community-mean species-level $d'_{plants}$ and $d'_{pollinators}$), separate GAMM/HGAMs were fitted for each predictor, with smooths estimated within levels of the potential methodological biases (sampling focus, habitat openness, region). Cells report the estimated degrees of freedom (edf) for the predictor's smooth in each level. edf = 0 indicates no effect (smooth shrunk to zero), edf ≈ 1 an approximately linear effect, and edf > 1 a non-linear effect. Significance refers to the predictor's smooth term: * p < 0.05, ** p < 0.01, *** p < 0.001. See Table S2 for variable definitions.

| Predictor | Sampling focus | | Habitat | | Region | |
|---|---|---|---|---|---|---|
| | Phytocentric | Zoocentric | Open | Forest | New World | Old World |
| **Complete dataset** | | | | | | |
| *$H_2'$* | | | | | | |
| latitude | 0.946*** | 0 | 0 | 4.266*** | 0.979 | 0 |
| MAT | 0.001** | 2.472** | 0.538*** | 4.553*** | 2.559*** | 4.592*** |
| SD_MMT | 0 | 0 | 4.875*** | 0 | 0.857** | 0 |
| MAP | 0 | 1.929* | 3.699** | 0 | 1.423** | 0 |
| SD_MMP | 0 | 1.051*** | 0.895** | 0 | 5.733*** | 0 |
| MAT×MAP | 0 | 5.301*** | 4.339*** | 0 | 8.494*** | 0 |
| mean NDVI | 0 | 0 | 0 | 0 | 3.32*** | 2.76*** |
| SD_NDVI | 0 | 1.48* | 1.51** | 0 | 0 | 1.77*** |
| Predicted plant species richness | 0 | 6.065*** | 0 | 0.644 | 4.103*** | 0 |
| Observed total species richness | 0 | 0 | 0 | 0 | 0.821* | 0 |
| Observed pollinator species richness | 0 | 1.987** | 0 | 0 | 4.676*** | 0 |
| Observed plant species richness | 0 | 0 | 0 | 0 | 0.881** | 0 |
| *$d'_{plants}$* | | | | | | |
| latitude | 0 | 2.089*** | 0* | 4.45** | 2.882*** | 2.097*** |
| MAT | 0 | 0.965*** | 0.623*** | 5.23*** | 0.526 | 0 |
| SD_ MMT | 1.371** | 0.572*** | 0 | 0 | 0 | 0 |
| MAP | 0.87** | 7.468*** | 0 | 0 | 0 | 0.375 |
| SD_MMP | 0 | 4.572*** | 2.892*** | 0 | 1.863* | 2.839*** |
| MAT×MAP | 0 | 1.905*** | 1.121* | 0 | 1.543** | 4.733*** |
| mean NDVI | 0 | 5.258*** | 0 | 2.039** | 0 | 3.447*** |
| SD_NDVI | 0*** | 4.684*** | 0 | 0 | 0.001 | 1.067 |
| Predicted plant species richness | 0** | 4.781*** | 4.877*** | 0** | 2.392*** | 1.225* |
| Observed total species richness | 0 | 2.838*** | 0 | 0 | 0.538 | 0 |

| | | | | | | |
|---|---|---|---|---|---|---|
| Observed pollinator species richness | 0 | 1.122*** | 0.133 | 0 | 4.433*** | 0 |
| Observed plant species richness | 0.81** | 7.115*** | 0.037** | 4.845*** | 0.004 | 0.355 |
| *$d'_{pollinators}$* | | | | | | |
| latitude | 0.006*** | 6.66*** | 8.527*** | 0*** | 5.873*** | 7.445*** |
| MAT | 0.995*** | 2.888*** | 0** | 7.313*** | 5.188*** | 5.295*** |
| SD_MMT | 0 | 0.975*** | 7.003*** | 0* | 3.504*** | 2.293*** |
| MAP | 0.319 | 3.991*** | 6.038*** | 0.001 | 5.926*** | 0.001 |
| SD_MMP | 0 | 4.673*** | 2.728*** | 0.891** | 0 | 8.022*** |
| MAT×MAP | 3.361** | 1.112** | 3.015*** | 20.399*** | 1.231*** | 7.05*** |
| mean NDVI | 0 | 7.194*** | 6.916*** | 7.369*** | 2.581*** | 0.028 |
| SD_NDVI | 0** | 6.604*** | 8.281*** | 4.164*** | 0.599*** | 3.055* |
| Observed total species richness | 0 | 3.37*** | 0 | 3.15*** | 0.95*** | 0 |
| Observed plant species richness | 0.004** | 6.702*** | 5.909*** | 0.001** | 7.282*** | 0.173* |
| **Full networks dataset** | | | | | | |
| *$H_2'$* | | | | | | |
| latitude | 0.342 | 0 | 4.712*** | 0 | 6.513*** | 0 |
| MAT | 0 | 0 | 0.006 | 0.636 | 2.826*** | 5.201*** |
| SD MMT | 0 | 0 | 5.107*** | 0 | 0.061 | 0 |
| MAP | 0* | 0 | 5.337*** | 0** | 6.278*** | 1.922* |
| SD_MMP | 0 | 0.039 | 0 | 0 | 4.859*** | 3.317* |
| MAT×MAP | 0 | 0 | 5.293*** | 1.118 | 8.437*** | 0.785 |
| mean NDVI | 0 | 0 | 0 | 1.374** | 5.438*** | 3.862*** |
| SD_NDVI | 0 | 0 | 0 | 0.228 | 0.001 | 1.969*** |
| Predicted plant species richness | 0 | 0 | 2.14** | 0 | 1.441** | 0 |
| Observed total species richness | 0 | 0 | 0 | 0 | 0.826* | 0 |
| Observed pollinator species richness | 0 | 0 | 0.326 | 4.153*** | 0 | 0 |
| Observed plant species richness | 0 | 0 | 0 | 1.762** | 0 | 0 |
| *$d'_{plants}$* | | | | | | |
| latitude | 0.691 | 0 | 2.076*** | 0 | 2.138*** | 0 |
| MAT | 2.257*** | 0 | 0 | 1.453** | 0.001* | 4.447*** |
| SD_MMT | 0 | 0 | 5.7*** | 4.014** | 0 | 0 |
| MAP | 0 | 0 | 3.212*** | 1.521*** | 5.209*** | 4.72*** |
| SD_MMP | 0 | 0 | 1.747** | 0.87** | 0 | 2.983*** |
| MAT×MAP | 0 | 0 | 0 | 1.03 | 1.692** | 6.791*** |
| mean NDVI | 0 | 0 | 0.831 | 0 | 2.838*** | 2.47*** |
| SD_NDVI | 0 | 0 | 0.671 | 0 | 0 | 1.776* |
| Predicted plant species richness | 2.897*** | 0 | 2.21*** | 0.001* | 0.568 | 0 |

| | | | | | | |
|---|---|---|---|---|---|---|
| Observed total species richness | 0 | 0 | 0 | 0 | 0.479 | 0 |
| Observed pollinator species richness | 0 | 0 | 0 | 0.24 | 0 | 0 |
| Observed plant species richness | 0.628 | 0 | 0.036 | 0 | 0 | 0 |
| *d'*$_{pollinators}$ | | | | | | |
| latitude | 0 | 0 | 0 | 0 | 6.667*** | 0.001*** |
| MAT | 0.002* | 0.579 | 4.149*** | 4.265*** | 0 | 3.948** |
| SD_MMT | 0.439 | 0 | 6.565*** | 4.741*** | 6.743*** | 0.002*** |
| MAP | 0.001* | 0 | 6.984*** | 1.727*** | 4.962*** | 0.001** |
| SD_MMP | 0.008*** | 1.628* | 0.001 | 0 | 7.215*** | 6.007*** |
| MAT×MAP | 0.766** | 0 | 15.203*** | 0.862* | 8.226*** | 3.238** |
| mean NDVI | 0 | 0 | 0 | 0 | 7.692*** | 7.52*** |
| SD_NDVI | 2.683*** | 0.489 | 7.721*** | 0 | 4.659*** | 0 |
| Observed species richness of functional groups | 0 | 0 | 0 | 0 | 0.86* | 3.48*** |
| Observed plant species richness | 0.114 | 0 | 1.159 | 0.001 | 2.159** | 0** |
| Functional-group networks | | | | | | |
| *$H_2$'* | | | | | | |
| latitude | 0 | 0 | 5.541*** | 4.643*** | 0.741*** | 5.004*** |
| MAT | 0** | 2.72** | 0 | 0.947*** | 0.006** | 5.869*** |
| SD_MMT | 0 | 0 | 6.916*** | 0 | 1.204*** | 0 |
| MAP | 0 | 0.782 | 0 | 0 | 2.12*** | 2.285*** |
| SD_MMP | 0.001** | 1.71** | 0 | 4.496*** | 3.397*** | 3.795*** |
| MAT×MAP | 0 | 2.751* | 3.583** | 0 | 1.82* | 5.899*** |
| mean NDVI | 0 | 0 | 0 | 0.771* | 3.065*** | 3.889*** |
| SD_NDVI | 0 | 0.803 | 0 | 1.117 | 0.971*** | 7.196*** |
| Observed species richness of functional groups | 0 | 0 | 0 | 0 | 1.19** | 0 |
| Observed plant species richness | 0.173 | 0 | 0.882 | 0 | 2.176*** | 0 |
| *d'*$_{pollinators}$ | | | | | | |
| latitude | 0.927*** | 0* | 5.391*** | 0** | 0.003*** | 7.08*** |
| MAT | 0* | 2.12*** | 0 | 5.895*** | 5.704*** | 5.54*** |
| SD_MMT | 0 | 0.924*** | 4.36*** | 4.738*** | 2.004*** | 0* |
| MAP | 0.511 | 2.446*** | 4.318*** | 0 | 6.872*** | 0.486* |
| SD_MMP | 0 | 1.781*** | 4.588*** | 3.968*** | 0.37* | 5.206*** |
| MAT×MAP | 0 | 3.888*** | 14.918*** | 4.158*** | 0.63 | 16.198*** |
| mean NDVI | 0** | 4.488*** | 4.154*** | 4.175*** | 6.297*** | 0.898*** |
| SD_NDVI | 0.172 | 0.912*** | 8.213*** | 0.002** | 0.367** | 2.353** |

| | | | | | | |
|---|---|---|---|---|---|---|
| Observed species richness of functional groups | 0 | 0.91** | 0 | 0 | 2.54*** | 0 |
| Observed plant species richness | 0 | 0 | 1.595*** | 0 | 0 | 0 |

**Table S7. Model comparisons identifying the best predictors of specialisation across datasets and metrics.** Results are shown for network-level *H₂′* and community-mean species-level $d'_{plants}$ and $d'_{pollinators}$ across the all networks, full networks, and functional-group networks datasets ($d'_{plants}$ not fitted for the functional-group dataset). Models were fitted within the GAMM/HGAM framework and compared by ΔAIC relative to the latitude-only baseline; rows are ordered by increasing ΔAIC within each dataset–metric block, with the best model highlighted in bold and marked with a star (★). "Covariates" lists potential methodological biases retained as random effects (sampling focus, habitat openness, region), selected based on their significance for each predictor model (see Table S6). $R^2adj$ gives variance explained by the whole model. See Table S2 for variable definitions and abbreviations, and Fig. 4 for smooth curves of the best-fitting predictors.

| Predictor | Covariates | $R^2_{adj}$ | Δ AIC |
|---|---|---|---|
| All networks dataset | | | |
| ***$H_2'$*** | | | |
| MAT ★ | habitat openness | 0.09 | -24.2 |
| SD_MMT | | 0.10 | -17.8 |
| latitude | region | 0.04 | 0.0 |
| MAP | region | 0.01 | 4.4 |
| SD_MMP | | 0.02 | 9.4 |
| MAP | | 0.01 | 10.3 |
| Observed pollinator species richness | | 0.01 | 13.0 |
| MAT×MAP | | 0.01 | 10.6 |
| Predicted plant species richness | | 0.00 | 17.6 |
| Observed plant species richness | | 0.01 | 17.7 |
| mean NDVI | | 0.00 | 17.8 |
| SD_NDVI | | 0.00 | 17.9 |
| Observed total species richness | | 0.09 | 24.2 |
| ***$d'_{plants}$*** | | | |
| SD_MMT★ | region, method | 0.23 | -8.3 |
| MAT | method | 0.19 | -5.1 |
| latitude | | 0.24 | 0.0 |
| Predicted plant richness | region, method | 0.17 | 13.1 |
| SD_NDVI | region, method | 0.17 | 19.7 |
| mean NDVI | method | 0.14 | 29.4 |
| MAP | method | 0.11 | 32.1 |
| SD_MMP | method | 0.13 | 36.9 |
| MAT×MAP | habitat openness, method | 0.11 | 41.2 |
| Observed species richness of functional groups | habitat openness, method | 0.10 | 44.0 |
| ***$d'_{pollinators}$*** | | | |
| MAT×MAP★ | region, method | 0.17 | 112.2 |
| MAP | region, method | 0.17 | 112.2 |
| Observed plant species richness | habitat openness, region, method | 0.10 | -85.6 |
| MAT | habitat openness, region, method | 0.12 | -31.4 |
| SD_MMP | habitat openness, region, method | 0.14 | -20.5 |
| SD_NDVI | habitat openness, region, method | 0.12 | -9.5 |
| latitude | habitat openness, region | 0.13 | 0.0 |
| SD_MMT | region, method | 0.12 | 6.8 |

| Predictor | Covariates | $R^2_{adj}$ | Δ AIC |
|---|---|---|---|
| Observed species richness of functional groups | habitat openness, region, method | 0.09 | 74.8 |
| mean NDVI | region, method | 0.10 | 103.7 |
| Full networks dataset | | | |
| ***H₂'*** | | | |
| MAT★ | habitat openness | 0.15 | -46.7 |
| MAT×MAP | | 0.16 | -34.7 |
| SD_MMT | habitat openness | 0.07 | -13.4 |
| mean NDVI | region | 0.09 | -12.3 |
| MAP | | 0.01 | -3.9 |
| SD_NDVI | region | 0.00 | 0.0 |
| latitude | region | 0.00 | 0.0 |
| Observed pollinator species richness | region | 0.00 | 0 |
| Observed plant species richness | region | 0.00 | 0 |
| SD_MMP | region | 0.0 | 0.0 |
| Observed total species richness | region | 0.00 | 5.3 |
| Predicted plant species richness | habitat openness | 0.15 | -46.7 |
| ***d'**$_{plants}$* | | | |
| MAT×MAP★ | habitat openness | 0.13 | -16.4 |
| Mean NDVI | | 0.07 | -14.8 |
| SD_NDVI | | 0.09 | -9.6 |
| MAT | | 0.04 | -3.4 |
| latitude | | 0.01 | 0.0 |
| MAP | | 0.01 | 3.0 |
| Predicted plant species richness | habitat openness | 0.03 | 3.1 |
| Observed plant species richness | | 0.00 | 3.9 |
| SD_MMP | | 0.00 | 4.3 |
| Observed pollinator species richness | | 0.00 | 4.6 |
| Observed total species richness | | 0.00 | 4.7 |
| SD_MMT | region | 0.01 | 5.4 |
| ***d'**$_{pollinators}$* | | | |
| MAT×MAP★ | region | 0.11 | -3.5 |
| latitude | habitat openness, region | 0.10 | -15.1 |
| SD_MMT | region | 0.09 | 0.0 |
| MAT | habitat openness, region | 0.09 | 5.3 |
| Observed plant species richness | habitat openness, region | 0.08 | 6.7 |
| mean NDVI | region | 0.07 | 7.4 |
| SD_NDVI | habitat openness, region | 0.06 | 20.9 |
| Observed species richness of functional groups | region | 0.06 | 43.7 |
| SD_MMP | | 0.07 | 71.6 |
| MAP | region | 0.04 | 68.0 |
| Functional-group networks | | | |
| ***H₂'*** | | | |
| latitude★ | region | 0.22 | 0.0 |
| MAP | region | 0.22 | 0.0 |

| Predictor | Covariates | $R^2_{adj}$ | Δ AIC |
| --- | --- | --- | --- |
| SD_MMT | region | 0.22 | 52.4 |
| SD_MMP | region | 0.21 | 38.2 |
| MAT×MAP | habitat openness | 0.20 | 58.1 |
| MAT | habitat openness | 0.12 | 120.8 |
| SD_NDVI | | 0.07 | 124.0 |
| mean NDVI | habitat openness | 0.06 | 133.8 |
| Observed plant species richness | habitat openness | 0.04 | 138.7 |
| Observed species richness of functional groups | | 0.12 | 149.3 |
| ***d'**pollinators*** | | | |
| MAT×MAP★ | region, habitat openness, method | 0.15 | -63.3 |
| latitude | habitat openness | 0.37 | -14.9 |
| MAP | habitat openness | 0.37 | -14.9 |
| SD_MMT | habitat openness, region | 0.34 | 0.0 |
| MAT | habitat openness | 0.25 | 120.9 |
| SD_NDVI | method | 0.21 | 166.5 |
| Observed plant species richness | habitat openness, region | 0.09 | 208.5 |
| mean NDVI | method | 0.10 | 257.9 |
| SD_NDVI | method | 0.06 | 269.2 |
| Observed species richness of functional groups | habitat openness, method | 0.14 | 279.9 |
